# Experimental structures of antibody/MHC-I complexes reveal details of epitopes overlooked by computational prediction

**DOI:** 10.1101/2023.12.01.569627

**Authors:** Lisa F. Boyd, Jiansheng Jiang, Javeed Ahmad, Kannan Natarajan, David H. Margulies

## Abstract

Monoclonal antibodies (mAb) to major histocompatibility complex class I (MHC-I) molecules have proved to be crucial reagents for tissue typing and fundamental studies of immune recognition. To augment our understanding of epitopic sites seen by a set of anti-MHC-I mAb, we determined X-ray crystal structures of four complexes of anti-MHC-I antigen-binding fragments (Fab) bound to peptide/MHC-I/β_2_m (pMHC-I). An anti-H2-D^d^ mAb, two anti-MHC-I α3 domain mAb, and an anti-β_2_-microglobulin (β_2_m) mAb bind pMHC-I at sites consistent with earlier mutational and functional experiments, and the structures explain allelomorph specificity. Comparison of the experimentally determined structures with computationally derived models using AlphaFold Multimer (AF-M) showed that although predictions of the individual pMHC-I heterodimers were quite acceptable, the computational models failed to properly identify the docking sites of the mAb on pMHC-I. The experimental and predicted structures provide insight into strengths and weaknesses of purely computational approaches and suggest areas that merit additional attention.

**ONE SENTENCE SUMMARY:** X-ray structures of anti-MHC-I/MHC-I complexes define epitopes overlooked by computational prediction.

## INTRODUCTION

The production and use of mAb have proved revolutionary for understanding fundamental issues of a range of problems including protein structure and function, cell differentiation, immunodiagnostics, and immunotherapy (*1–3*). MAb directed against the cell surface histocompatibility antigens H2 in the mouse and HLA in the human have been particularly useful in a wide range of applications (*4–7*), including the exploration of structural aspects of MHC molecules (*8, 9*) and HLA typing for organ transplantation (*10*). Anti-MHC-I mAb have also been used for biosynthetic studies of the cell biology and assembly of MHC molecules as they proceed from synthesis in the endoplasmic reticulum (ER) through steps of folding, glycosylation, peptide acquisition, and trafficking to the cell surface for recognition by receptors on T cells, NK cells, and other immune and inflammatory cells (*9, 11, 12*). Recently, in efforts to target specific peptide/MHC (pMHC) complexes characteristic of tumor cells, mAb that mimic T cell receptor (TCR)-mediated recognition have been isolated and are in development as part of the therapeutic armamentarium for a variety of malignancies (*13*). TCR mimic (TCRm) mAb have potential value in treating autoimmune diseases as well (*14*).

Many anti-MHC-I mAb were initially defined by their reactivity against genetically well-defined strains of inbred animals (*15*) or characterized by examination of reactivity against panels of well-known cell lines or purified HLA molecules (*16*). Some anti-MHC-I mAb have been studied further for their recognition of the cell surface products of *in vitro* manipulated MHC-encoding genes to map the domain location of specific epitopes (*17–20*) in order to identify focal amino acid residues that define antibody recognition sites (*21–24*). These anti-MHC-I mAb have contributed to an understanding of specific sites that define MHC polymorphism and control interaction with TCR, coreceptors, NK receptors (NKR), or other inhibitory or activating receptors (*25–27*). Although such approaches have been valuable for a broad identification of the MHC epitopes recognized by such mAb, structural analysis offers to elucidate further details of the binding sites and to provide insight into which mAb may compete for binding by ligands with known sites of interaction. These structures can provide a basis for engineering antibodies with increased affinity or improved specificity. In addition, precise knowledge of the antigenic epitopic residues provides a structural basis for the transfer of specific recognition sites to other allelomorphs or even unique engineered proteins.

To understand better the details of anti-MHC-I mAb, we determined experimentally by X-ray crystallography the structures of complexes of mAb Fab with pMHC-I. We crystallized complexes of four anti-murine MHC-I mAbs: two that bind distinct regions of H2-D^d^ (34-5-8 (α2 domain) and 34-2-12 (α3 domain) (*28*)), one that binds the conserved α3 domain of both H2-L^d^ and -D^b^ (28-14-8 (*29*)), and one that discriminates a single amino acid polymorphism of the β_2_m light chain subunit of MHC-I (S19.8 (*30, 31*)). These experimental structures reveal details of the footprints of their respective Fab on pMHC-I consistent with previous biochemical, genetic, and immunological studies. In addition, the structures pinpoint side chain interactions, explain allele specificity, and shed light on conformationally plastic regions of pMHC, particularly with respect to changes observed in the α2 domain on peptide binding.

With the X-ray structures in hand, we evaluated the ability of one computational algorithm, AlphaFold-Multimer (AF-M) (*32*) to predict and visualize complexes of these selected antibodies with their respective MHC-I protein antigens. Deep-learning methods, such as AlphaFold, have been remarkably successful for prediction of protein structures from amino acid sequence (*33, 34*), particularly with respect to individual domains of structured proteins. The release of three-dimensional models of the entire human proteome already promises rapid progress in rational approaches to drug discovery and understanding fundamental mechanisms of cellular biochemistry (*35*). Although computational determination of the organization of multidomain proteins and multimolecular complexes is clearly a more challenging problem than domain prediction alone (*36*), AF-M (*32*) offers an opportunity for predicting and evaluating higher order interactions (*37*). The availability of AF-M implemented in ChimeraX (*38, 39*) and linked to Google Colab servers (*40*) permits rapid assessment of models of a variety of protein complex structures. We applied this modeling approach to these four mAb/MHC-I complexes. Although the resulting computation generated good models for the previously well-known MHC-I/β_2_m complexes, the models revealed shortcomings in prediction of Fab and variable region fragment (Fv) structures. Computational models failed to properly identify the sites where the Fv V_H_V_L_ docked on the MHC-I. The discrepancies between experiment and prediction arise from difficulties in establishing proper domain relationships as shown by elbow angle, ambiguities in loop structures, particularly the Ab complementarity determining regions (CDRs), and to complexities in the docking of Ab with protein antigens. Accumulation of experimental structural data on protein antigen/Fab complexes should provide a more extensive database for improvement of algorithms for structure prediction.

## RESULTS

### X-ray Structures of mAb bound to MHC-I molecules

To understand details of the interactions of the four anti-MHC-I mAb, we purified refolded recombinant H2-D^d^/ and H2-D^b^/β_2_m complexes containing high affinity peptides and prepared Fab of the mAb as described in detail in the Methods. Diffraction quality crystals of Fab/MHC-complexes were obtained, and X-ray data sets were collected at resolutions from 2.60 to 2.90 Å (see Table 1). Complexes of the four Fab/MHC-I complexes were readily solved by molecular replacement and refined as summarized in Materials and Methods and Table 1. Previous serological and functional studies had mapped 34-5-8 to the α1α2 peptide-binding domain and 34-2-12 and 28-14-8 to the α3 domains of H2-D^d^ and H2-L^d^/D^b^ respectively (*17*). Detailed mutational analysis further localized the 34-5-8 epitope (peptide-dependent, but not peptide specific (*41*)) to particular residues of the α2 domain (*42–44*). S19.8 recognized MHC-I molecules containing the β_2_m^b^ allelomorph (Ala85) and not β_2_m^a^ (Asp85) (*31*).

**Table 1.** X-ray data collection and refinement statistics.

|  | **Fab34.5.8/**<br>**H2-D<sup>d</sup>** | **Fab34.2.12/**<br>**H2-D<sup>d</sup>** | **Fab28.14.8/**<br>**H2-D<sup>b</sup>** | **FabS19/**<br>**H2-D<sup>d</sup>** |
| --- | --- | --- | --- | --- |
| **PDBID** | **8TQ8** | **8TQ7** | **8TQA** | **8TQ9** |
| **Data collection** |  |  |  |  |
| Space group | P2 | P2<sub>1</sub>2<sub>1</sub>2<sub>1</sub> | P2<sub>1</sub> | I222 |
| Cell dimensions |  |  |  |  |
| *a*, *b*, *c* (Å) | 87.09, 50.89, 224.83 | 87.59, 113.33, 196.37 | 71.39, 70.19, 94.34, | 54.90, 183.18, 216.72 |
| $\alpha$, $\beta$, $\gamma$ (°) | 90.0, 90.0, 90.0 | 90.0, 90.0, 90.0 | 90.0, 98.0, 90.0 | 90.0, 90.0, 90.0 |
| Resolution (Å)* | 81.21-2.69<br>(2.79-2.69) | 56.62-2.80<br>(2.90-2.80) | 52.94-2.60<br>(2.69-2.60) | 54.18-2.90<br>(3.00-2.90) |
| *R*<sub>sym</sub> or *R*<sub>merge</sub> | 0.145 (0.927) | 0.099 (1.316) | 0.067 (0.479) | 0.108 (1.158) |
| *I*/$\sigma$(*I*) | 7.6 (1.3) | 12.9 (1.2) | 13.1 (2.6) | 14.4 (1.8) |
| Completeness (%) | 97.9 (99.3) | 96.9 (93.5) | 98.8 (97.7) | 95.4 (89.8) |
| Redundancy | 3.8 (3.9) | 6.7 (6.7) | 3.8 (3.9) | 7.4 (7.2) |
| *R*<sub>pim</sub> | 0.085 (0.543) | 0.041 (0.547) | 0.039 (0.276) | 0.042 (0.455) |
| CC<sub>1/2</sub> | 0.990 (0.560) | 0.998 (0.528) | 0.997 (0.855) | 0.998 (0.648) |
| Twin | (h,-k,-l) | none | none | none |
| **Refinement** |  |  |  |  |
| Resolution (Å)* | 81.21-2.69<br>(2.79-2.69) | 56.62-2.80<br>(2.90-2.80) | 52.94-2.60<br>(2.69-2.60) | 54.18-2.90<br>(3.00-2.90) |
| No. unique reflections<sup>§</sup> | 54123 (5199) | 48825 (4505) | 28375 (2837) | 24406 (2174) |
| *R*<sub>work</sub>/*R*<sub>free</sub> (%) | 21.4/25.3<br>(31.1/34.6) | 22.4/27.8<br>(33.0/36.8) | 20.3/26.5<br>(27.6/38.1) | 20.6/24.9<br>(30.7/37.2) |
| No. atoms | 12707 | 12724 | 6431 | 6385 |
| Protein | 12446 | 12584 | 6302 | 6315 |
| Water/ligands | 255/6 | 134/6 | 67/62 | 59/11 |
| B-factor Wilson/Average | 57.8/65.3 | 55.2/73.2 | 37.7/44.8 | 59.0/62.6 |
| Protein | 65.8 | 73.5 | 44.8 | 62.7 |
| Water/ligands | 39.8/46.0 | 38.66/168.3 | 34.9/60.6 | 41.8/111.0 |
| R.m.s. deviations |  |  |  |  |
| bond length (Å) | 0.003 | 0.005 | 0.002 | 0.004 |
| bond angle (°) | 0.62 | 0.79 | 0.50 | 0.69 |
| Ramachandran |  |  |  |  |
| favored (%) | 88.6 | 94.8 | 93.7 | 93.8 |
| allowed (%) | 9.5 | 4.5 | 4.9 | 5.7 |
| outliers (%) | 1.9 | 0.8 | 1.4 | 0.5 |
| Molprobability |  |  |  |  |
| clashscore | 12.8 | 10.7 | 5.1 | 7.5 |

(\*Values in parenthesis are for highest resolution shell, <sup>§</sup> Values in parenthesis are the number of reflections for *R*<sub>free</sub>)

### Structure of Fab34-5-8 bound to H2-D^d^

As shown in Fig. 1 and Supplementary Table 1, 34-5-8 interacts only with residues of the H2-D^d^ α2 domain, exploiting contacts of its V_H_ and V_L_. The overall structure of the pMHC-I complex is the same as that of some 19 previously determined H2-D^d^ structures in the protein data bank (PDB) and reveals RMSD values for H2-D^d^ ranging from 0.762 to 1.545. Similarly, the RMSD of β_2_m varies from 0.434 to 0.824 Å. The basic folds of the Fab V_H_V_L_ and C_H1_C_L_ of 34-5-8 are clearly representative of a host of previously determined Fab structures. The footprint of the Fab V_H_V_L_ on H2-D^d^ is determined by interactions of 19 residues of the α2 domain (but none of β_2_m), including 17 that contact the Fab H chain and 7 that contact the L chain of which 5 are in common. The buried surface area of the MHC-I chain is 925 Å^2^, of which 714 Å^2^ are due to the Fab H chain, and 211 Å^2^ to the L chain (see Supplementary Table 2), and the shape complementarity (*S_c_* (*45*)) between V_H_V_L_ and H2-D^d^ is 0.637, characteristic of antigen/Ab interfaces. The region of the footprint is illustrated in Fig. 1D with contacts of the Fab to residues of the β-sheet floor (residues 104-111 and 127-132) and to several of the α2-2 helix (154, 157, 161, 162, 165 and 169). Residues 104-108 form a tight turn (focused on E104 and R106) that is engaged by H chain residues of CDRH1 (A28, S31, and Y32), CDRH2 (residues 52-57) and CDRH3 (residues 99-104) (Fig. 1E and 1F). L chain residues of CDRL2 (Y52, R54, N57, D59, S60) and of CDRL3 (E97 and W101) also contact H2-D^d^.

**Fig. 1.**
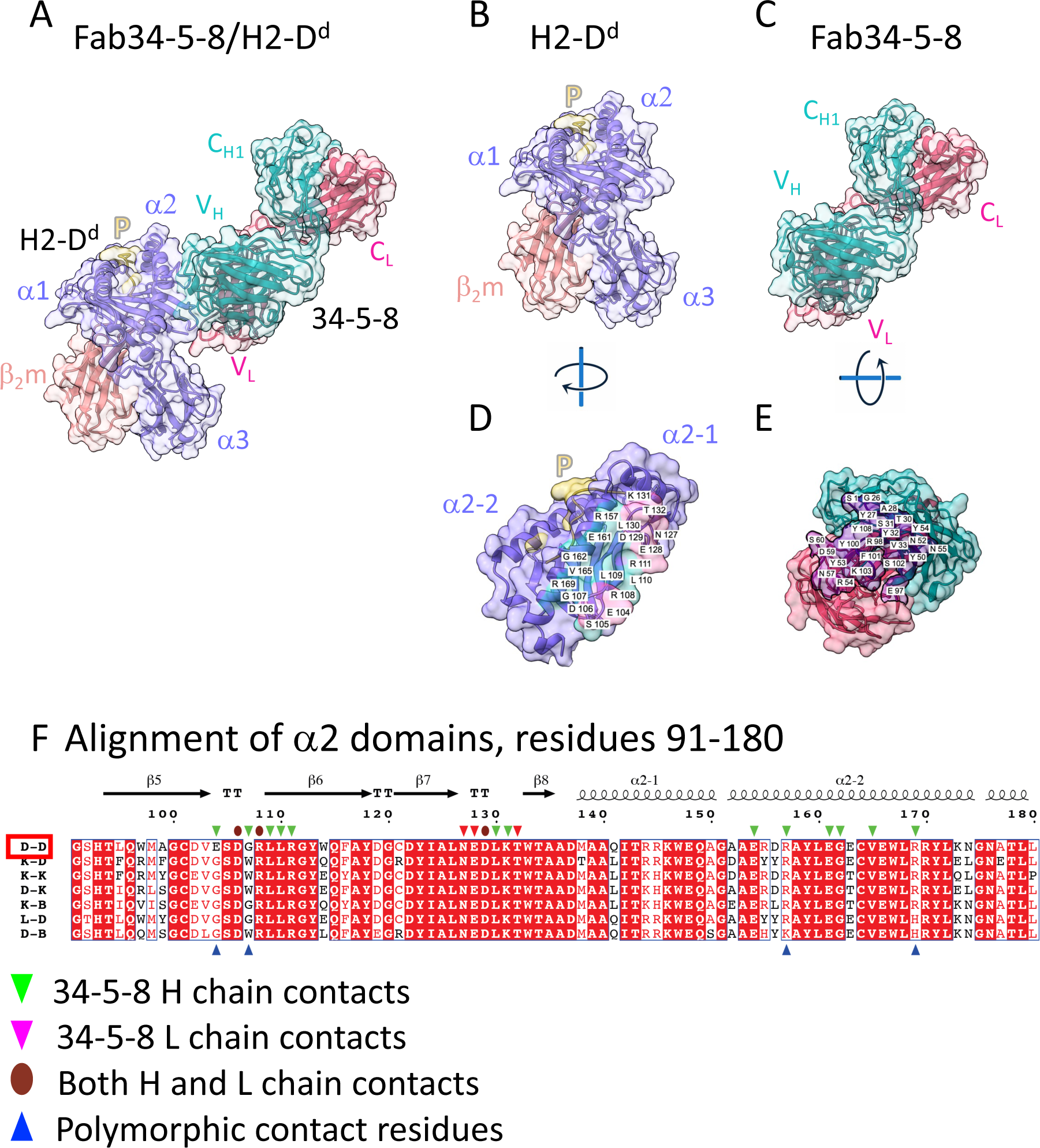
X-ray structure of complex between Fab34-5-8 and H2-D^d^ reveals footprint on a2 domain. Illustration in cartoon and partially transparent surface representation of (A) complex of Fab34-5-8 with H2-D^d^. H2-D^d^ a (Heavy) chain, medium slate blue, β_2_m (light coral), peptide (yellow), Fab H chain, dark cyan, Fab L chain, red; (PDB 8TQ8); (B) pMHC complex alone; (C) Fab34-5-8 H/L alone; (D) H2-D^d^ α1α2 and peptide rotated to allow visualization of interface; (E) Fab34-5-8 combining site rotated to permit visualization of H (cyan) and L (red) contact residues. (F) structure guided alignment of the α2 domains of the indicated mouse MHC-I molecules. Secondary structure elements above the alignment determined from PDB 3ECB. Contacts of Fab H and L chains are indicated (downward cyan (H) and magenta (L) arrows and brown (both) ovals), and residues that both contact the Fab and are polymorphic are indicated by upward blue arrows. Alignment was performed with Clustal Omega https://www.ebi.ac.uk/Tools/msa/clustalo/ (*91*) and illustration prepared with ESPript 3.0 https://espript.ibcp.fr/ESPript/cgi-bin/ESPript.cgi (*92*).

Examination of the Fab/H2-D^d^ interface explains the private specificity of the mAb. The unique H2-D^d^ residue E104 as well as G107 and R157 are contacts made by the H chain, while the L chain contacts residues conserved among a sampling of other murine MHC-I allelomorphs (see Fig. 1F). Previous mutational analysis revealed reduced 34-5-8 reactivity of E104G and G107W, and the suggestion of improved binding of W97R (*46*), a residue that sits in the peptide binding groove but does not make direct contact with mAb 34-5-8. Since the W97 side chain is directed into the peptide binding groove and is distant (W97 NE1 to H chain Y100 O is 16.8 Å away) from the site of interaction with the 34-5-8 Fab, this likely reflects a change in β-strand 5 that influences the conformation of the 103 to 109 loop which serves as a direct site for interaction with the Fab H and L chains. Another region of H2-D^d^ that contacts Fab 34-5-8, residues 127-132 (see Fig. 1F) also plays a crucial role in recognition by this mAb, as evidenced by the observation that the H2-D^dm6^ mutation W133R obliterates binding to 34-5-8 (*47*). Although the completely conserved W133 is adjacent to but not directly in contact with the mAb, our modeling of the R substitution at this position clearly indicates distortion of the β8 strand and the peptide binding groove. Thus, the X-ray structure of the Fab34-5-8/H2-D^d^ complex confirms the results of earlier exon-shuffling and mutagenic studies and explains in detail the specificity of the mAb. Furthermore, the structure indicates that the peptide-dependent, but not peptide specific, recognition by the mAb reflects the sensitivity of the 104-111 loop as an indicator of peptide binding.

### Structure of Fab34-2-12 bound to H2-D^d^

The Fab34-2-12/H2-D^d^ complex, illustrated in Fig. 2, shows that the Fab H and L chains recognize three loops at the membrane proximal surface of the H2-D^d^ α3 domain. The p/H2-D^d^/β_2_m structure closely resembles that of other independently solved H2-D^d^ structures (MHC-I heavy plus light, RMSD from 0.708 to 2.163; heavy chain RMSD from 0.692 to 1.775, and β_2_m from 0.289 to 0.887 Å). The interface between the Fab and H2-D^d^ involves 21 residues of the H2-D^d^ heavy chain, 19 of which interact with the Fab H chain and two with the Fab L chain, burying some 757 Å^2^ of the H2-D^d^ α3 domain (Supplementary Tables 1 and 2) with an *S_c_* value (*45*) of 0.668. Of particular note is that 34-2-12 envelopes the region of the α3 domain, residues 219 to 227 (Fig. 2F), a region that is bound by the costimulatory T cell molecule, CD8αβ, described crystallographically by PDB 3DMM (*48*). Additionally, Fab34-2-12 binds membrane proximal loops of H2-D^d^ (residues 194-197 and 247-257), suggesting that it might bind H2-D^d^ molecules lying in a supine position on the cell surface–the carboxyl-terminal strand from β17 would be expected to have access to a peptide strand connecting this to the transmembrane region of H2-D^d^ which would run from 271 to about residue 288. (Evidence for such an orientation of H2-K^b^ on a lipid bilayer has been reported (*49*)). The observation that 34-2-12 binds only amino acid residues of the α3 domain is consistent with experiments showing that it blocks function of CD8^+^ cytolytic cells (*21*), and stains cells expressing recombinant truncated H2-D^d^ α1α2 deletion mutants (*50*) and MHC-II/MHC-I hybrid molecules (*51*). H2-D^d^ mutants that abrogate 34-2-12 binding also reveal diminished susceptibility to cytolysis by CD8^+^ cells (*21, 52*).

**Fig. 2.**
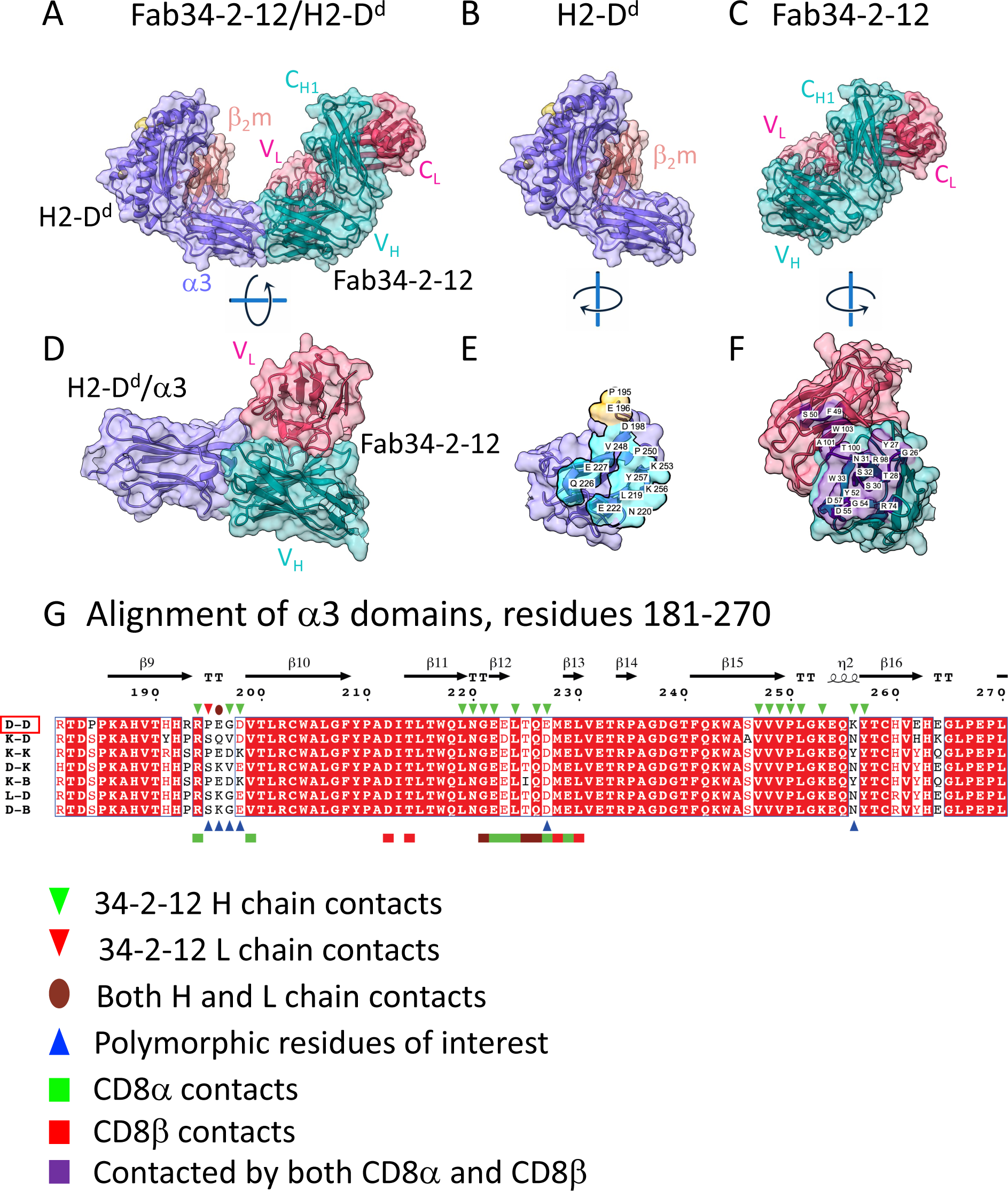
X-ray structure of complex between Fab34-2-12 and H2-D^d^ reveals footprint on α3 domain. Illustration in cartoon and partially transparent surface representation of (A) complex of Fab34-2-12 with H2-D^d^ (Colors of indicated chains as in Fig. 1); (B) pMHC (H2-D^d^) complex alone; (C) Fab34-2-12 H/L alone; (D) H2-D^d^ α3 alone with V_H_V_L_ rotated to visualize contacts; (E) pMHC rotated to allow visualization of Fab contacts; (F) Fab rotated to allow visualization of contacts to pMHC; (G) alignment of amino acid sequences of α3 domains of the indicated murine MHC-I molecules, performed as described in legend to Fig. 1. Contacts of Fab34-2-12 H and L chains are indicated, as are contacts to CD8α and CD8β as determined in PDB 3DMM (*48*).

### Structure of Fab28-14-8 bound to H2-D^b^/β_2_m^b^

Consistent with previous mapping, the structure of the Fab28-14-8/H2-D^b^ complex, illustrated in Fig. 3, reveals direct contact of both the H and L chains primarily with residues of the α3 domain of H2-D^b^. Additionally, two residues of the α2 domain (conserved R111 and E128) and also one residue of the β_2_m light chain (conserved D59) are bound by the Fab. The Fab buries 852 Å^2^ of the H2-D^b^ heavy chain and rather little (11 Å^2^) of the β_2_m light chain (Supplementary Table 2), and the *S_c_* of the interface (*45*) is 0.704. The focus of the Fab is on an extended region in the center of the α3 domain, involving residues 212-226, a region that largely overlaps the main contacts of the CD8αβ heterodimer with H2-D^d^ (PDB 3DMM) (*48*). The minimal contact with β_2_m is consistent with earlier findings that 28-14-8 binds β_2_m-free surface MHC-I molecules as well as H2-D^b^ expressed in a β_2_m negative cell line and α1α2 deletion mutants of H2-D^b^ as well (*53*). Structural alignment of H2 allelomorph sequences with attention to residues that distinguish those molecules that bind 28-14-8 (H2-D^b^, -L^d^) from those that do not (H2-K^b^,-D^d^,-K^d^,-K^k^,-D^k^) indicates that R260, bound by Fab H chain Y101 and L chain G91 is critical for the allelic specificity of the Ab.

**Fig. 3.**
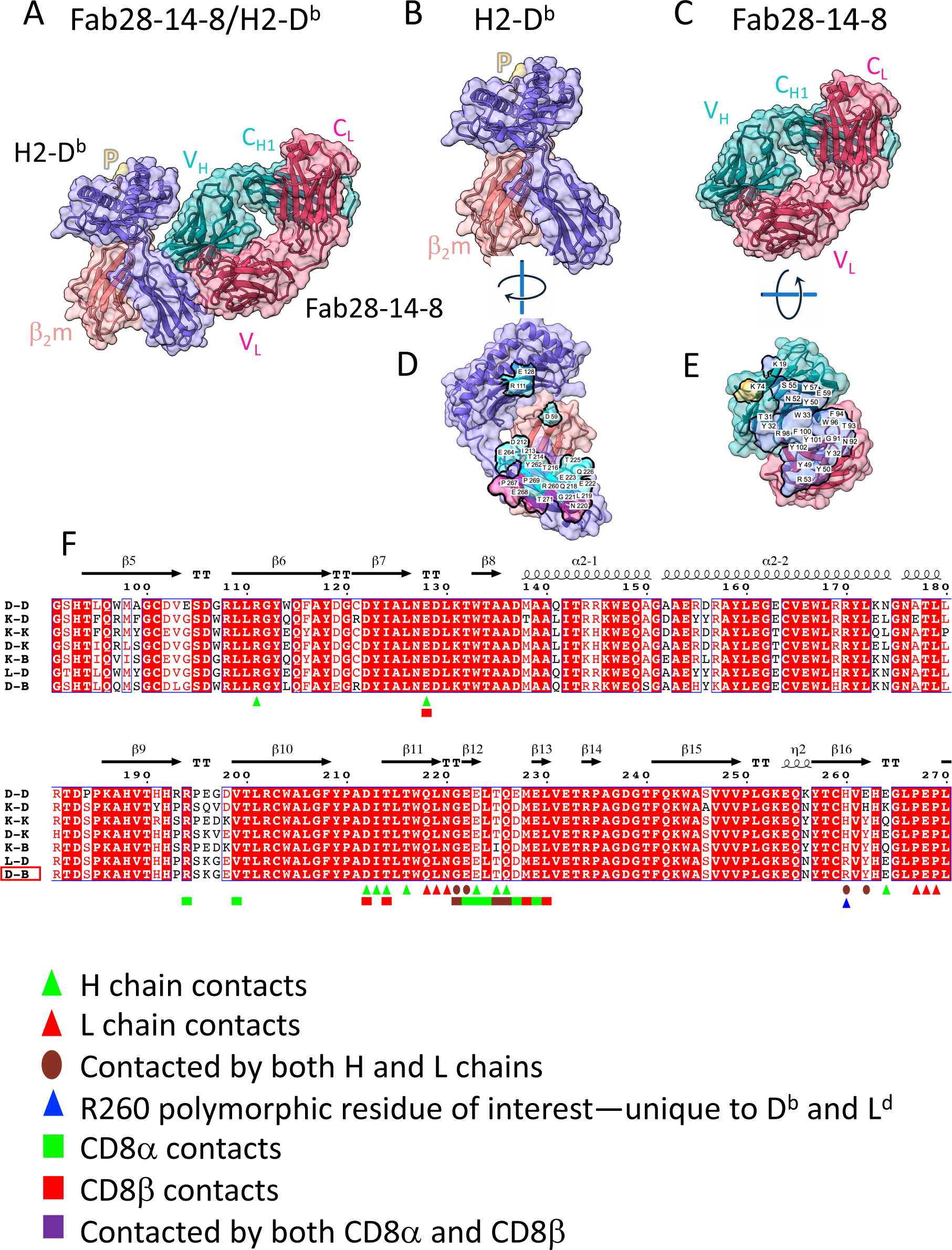
X-ray structure of complex between Fab28-14-8 and H2-D^b^ reveals footprint on α2 and α3 domains. Illustration in cartoon and partially transparent surface representation of (A) Fab28-14-8 with H2-D^b^ (Colors of indicated chains as in Fig. 1); (B) pMHC complex alone; (C) Fab28-14-8 H/L alone; (D) H2-D^d^ rotated to visualize contacts to Fab; (E) Fab rotated to allow visualization of contacts to pMHC; (F) alignment of amino acid sequences of α2 and α3 domains of the indicated murine MHC-I molecules, performed as described in legend to Fig. 1. Contacts of Fab28-14-8 H and L chains are indicated, as are contacts of H2-D^d^ to CD8α and CD8β as determined in PDB 3DMM (*48*).

### Structure of FabS19.8 bound to H2-D^d^/β_2_m^b^

Fab S19.8 represents a specificity focused on the β_2_m light chain of H2 complexes rather than on polymorphic structures of the H2 heavy chains. The S19.8 hybridoma (SJL (H2^s^/β_2_m^a^) anti-B10.S (H2^s^/β_2_m^b^)) was originally designated anti-ly-m11.2 (*30*) and subsequently identified as anti-β_2_m^b^ (*31, 54, 55*). Binding studies using purified S19.8 and a selection of recombinant mouse and human MHC-I molecules prepared with both allelic forms of β_2_m indicate that S19.8 distinguishes β_2_m^b^ from β_2_m^a^ when in complex with H2-K^b^ (Supplemental Fig. 1). Additionally, the mAb binds free β_2_m^b^ but does not bind human β_2_m. The *K*_D_ value for binding to H2-K^b^/β_2_m^b^ is 1.3 × 10^-6^ M while that for β_2_m^b^ alone is approximately 6-fold weaker at 8.3 × 10^-6^ M (Supplemental Fig. 1). The X-ray structure of the FabS19.8/H2-D^d^/ β_2_m^b^ complex (Fig. 4) reveals the detailed explanation for this binding behavior. S19.8 H and L chains engage 1097 Å^2^ of the H2-D^d^/β_2_m^b^ complex, with the greater area contributed by β_2_m, 906 Å^2^ vs. 191 Å^2^ of the H2-D^d^ heavy chain (see Supplementary Table 2). The overall disposition of the S19.8 Fab is toward β_2_m (18 residues (Fig. 4 G)) and six residues of H2-D^d^ (α1 residues S13, R14, P15, F17 and α2 G91, S92—Fig. 4F)(see Supplementary Table 1). β_2_m residue A85 defines the polymorphism of β_2_m^b^ which is at the center of the S19.8 H and L chain contact area. Thus, the β_2_m^a^ chain, with D at 85 would be expected to be sterically incompatible with the S19.8 interaction (see Fig. 4 H, I). Reactivity of S19.8 with rat β_2_m has been reported and is consistent with the identity of 13 contact residues (1, 2, 4, 32, 35, 36, 45, 81, 82, 83, 84, 89, 90) and the similarity of two (85, 87). Of note are differences at positions 34, 38, and 88 that may be sufficient to affect the affinity of the interaction. The decreased reactivity of MHC-I complexes containing human β_2_m with S19.8 (Supplementary Fig. 1) likely reflects major differences at positions 34 (human D for H), 38 (human D for Q) and 89 (human Q for E) (see Fig. 4G). Further binding, mutagenic, and structural studies will be needed to reveal the details of the contributions of each of the residues of the S19.8/β_2_m interface in different species. The ability of S19.8 to distinguish a single amino acid substitution at the center of its interface, despite a rather modest affinity, is a striking example of how low affinity interactions may be the basis of clearcut molecular discrimination.

**Fig. 4.**
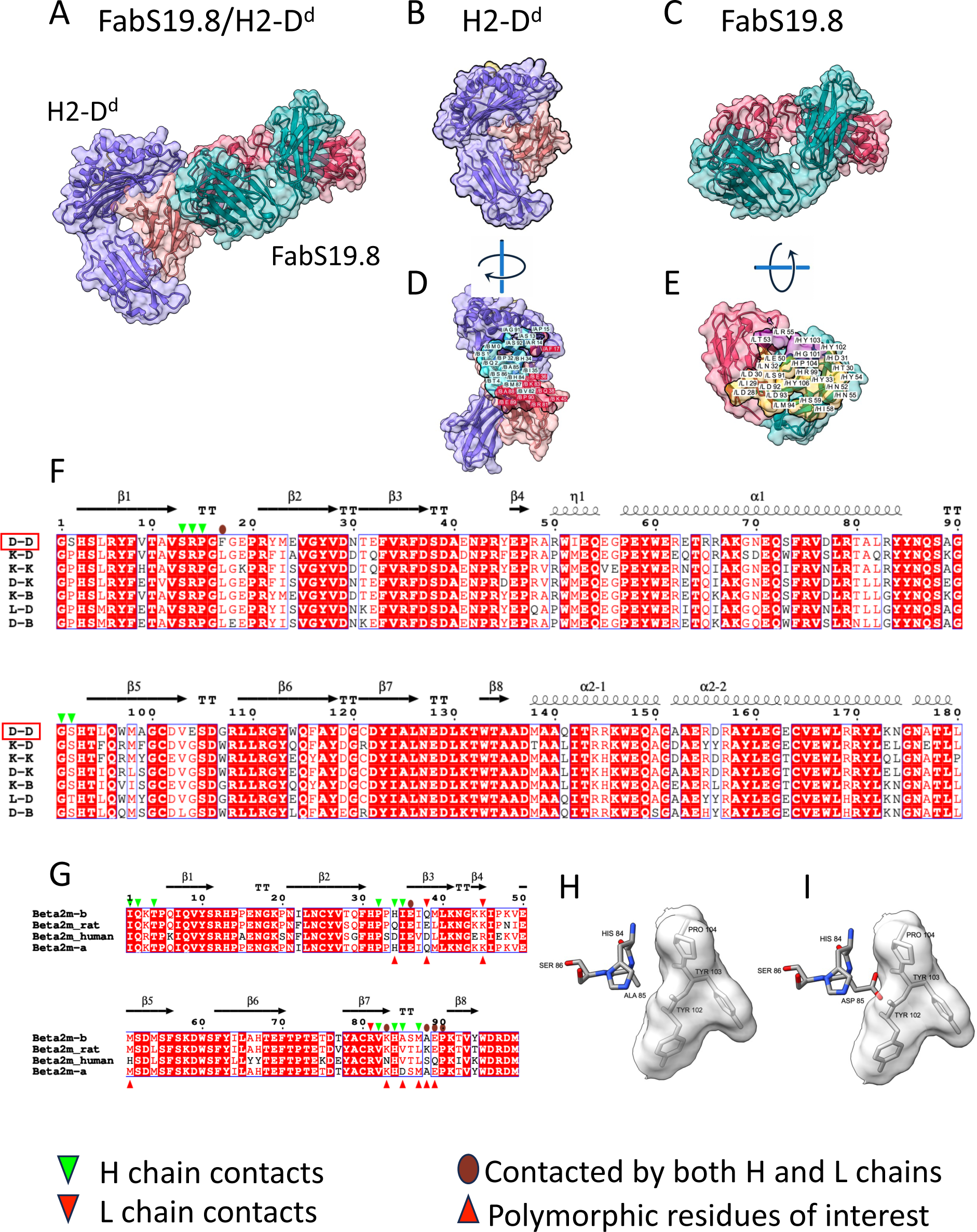
X-ray structure of complex between FabS19.8 and H2-D^d^/β_2_m^b^ reveals footprints on α1 and α2 domains as well as on β_2_m. Illustration in cartoon and partially transparent surface representation of (A) complex of FabS19.8 and H2-D^d^/β_2_m^b^ (Colors of indicated chains as in Fig. 1); (B) pMHC complex alone; (C) FabS19.8 H/L alone; (D) H2-D^d^/β_2_m^b^ rotated to visualize contacts to Fab; (E) Fab rotated to allow visualization of contacts to pMHC; (F) alignment of amino acid sequences of α1 and α2 domains of the indicated murine MHC-I molecules; (G) alignment of the indicated β_2_m sequences; (H) close-up illustration of sticks representation of β_2_m^b^ addressing surface of S19.8 H chain residues 102-104; (I) model of clash encountered by substitution of β_2_m Ala85 to Asp, superposed on structures illustrated in (H).

### AlphaFold-Multimer Models of mAb/MHC-I complexes

Having solved the structures of these four Fab/MHC-I complexes, we reasoned that they might provide a fair test of the ability of AF-M (*32*) to predict the structure of each of the Fab/MHC-complexes determined experimentally. Thus, the amino acid sequences of the component chains of each of the four complexes were submitted for prediction as described in Materials and Methods. In general, the predicted individual models of the basic MHC-I and β_2_m folds were good, and pMHC-I trimer or V_H_V_L_ and C_H1_C_L_ heterodimers were reasonable (as indicated by predicted aligned error (PAE) plots in AlphaFold). The pMHC-I heterotrimers were compared to the AF-M predictions as summarized in Supplementary Fig. 2. Fab34-5-8/H2-D^d^ complexes showed good agreement of the determined vs. predicted pMHC-I (overall all-atom RMSD of 1.814 Å and MHC-I H chain alone of 1.330 Å). For the β_2_m, however, a larger RMSD (2.690 Å) was noted (Supplementary Fig. 2A, B), indicative of some strand differences, and the peptide model also showed some distortion, evidenced by an RMSD of 2.574 Å. Differences among the α1α2 peptide binding domain (RMSD of 1.259 Å) and the α3 domain independently were smaller (1.456 Å). Evaluation of the pMHC-I structures and models of the other three complexes showed several individual variations. The H2-D^b^ complex with Fab28-14-8 revealed a distortion of the MHC-I H chain 103-110 loop (RMSD 5.428 Å), a region not contacted by the Fab (Supplementary Fig. 2A,C). Analysis of the H2-D^d^ complex with S19.8 showed that although the MHC-I H chain and β_2_m were rather similar between structure and AF-M model, loops of the peptide and α1 domain (residues 13 to 20) differed.

For the complete four domain Fabs, we examined the differences between the variation in elbow angles describing the relationship of V_L_V_H_ to C_L_C_H1_ which varied widely. Differences in the elbow angles of the experimental structures as compared with those of AF-M models are summarized in Supplementary Table 3. Thus, the X-ray determined elbow angle for Fab34-5-8 is 150° as compared with the AF-M model of 136°. Similarly, Fab34-2-12, Fab28-14-8, and FabS19.8 differ by 1.0, 28.0 and 16.0° respectively. Additional comparisons of the X-ray vs. AF-M models were carried out by calculating all atom RMSD which varied widely (Supplementary Fig. 3). Clearly, wide variation exists among the residues of the V region of the Fab H chains, particularly in the CDR loops (Supplementary Fig. 4), with 34-2-12 displaying the least variation and 28-14-8 showing the most. The final step, the prediction of the docking site of the Fab onto the MHC-I molecules, did not agree with the experimentally determined structures and was grossly incorrect for all four Fab/MHC-I complexes. This is evaluated graphically in Fig. 5, and quantitatively, using DockQ (*56*) in Supplementary Table 4.

**Fig. 5.**
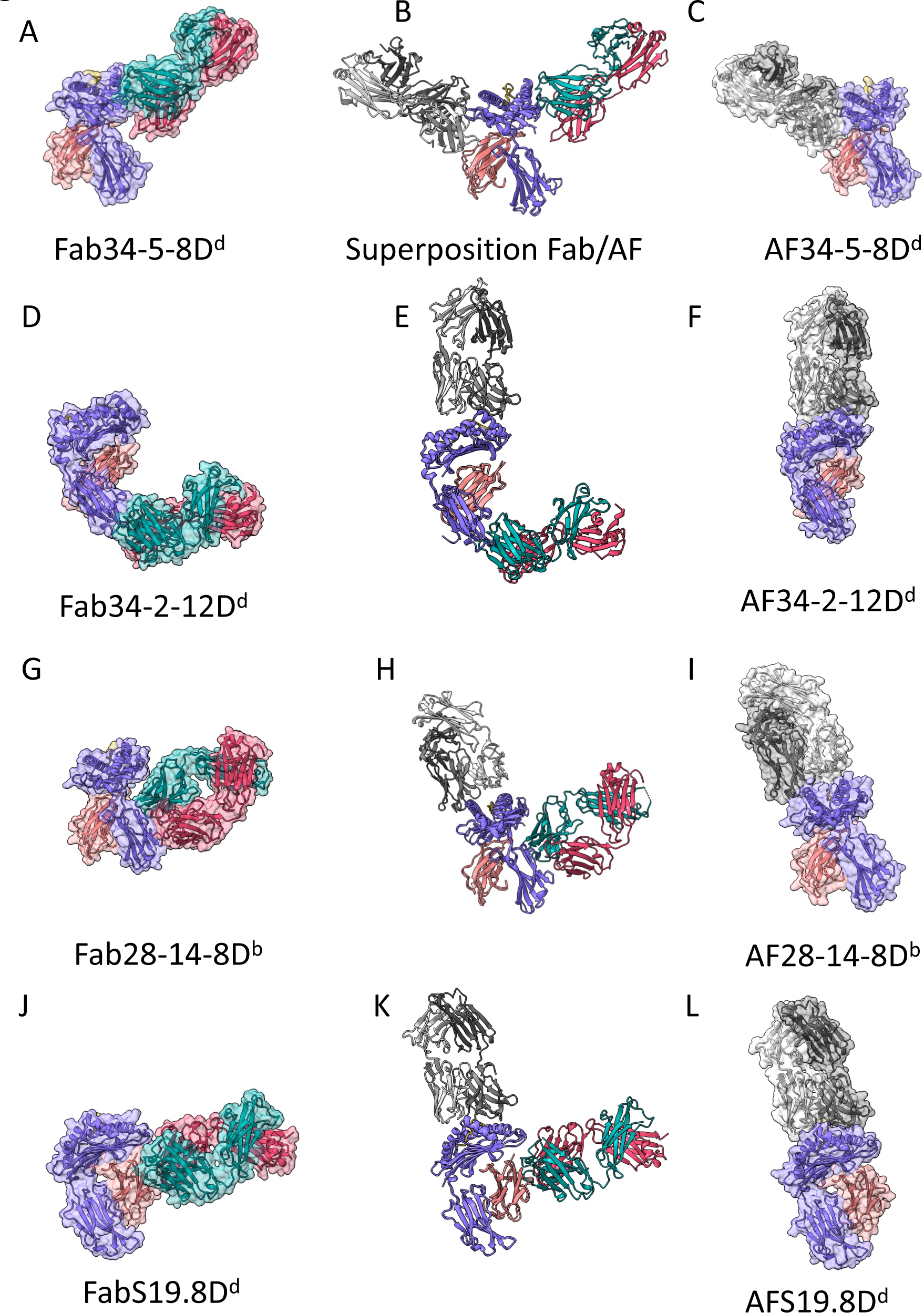
Comparison of X-ray structures of Fab/MHC complexes to AF-M models reveals differences in experimental vs. computed docking sites of the Fab. AF-M predictions of Fab/MHC complexes were accomplished as described in Materials and Methods, and the X-ray determined models were compared to the “best_model” from each of the AF-M predictions. The indicated MHC-I heavy chains of the structures were superposed and the complete models were displayed in ChimeraX. Panels (A, D, G, J) show the X-ray structures and panels C, F, I, and J the AF-M models, while panels B, E, H, and K show the superpositions. MHC H chain is purple and L chain coral, X-ray Fab H chain is cyan and L chain red. For the AF-M models MHC colors are the same, but Fab H chain is dark grey and L chain is light grey. For the superposed Fab/MHC complex structures and predictions, we calculated the difference in the location of the center of mass (COM) of each Fab H and L chain. For 34-5-8 (B), H chain differed by 86.2 Å and L chain by 100.0 Å). Similarly, for 34-2-12 (E), H and L differed by 95.1 and 112.9 Å, respectively; for 28-14-8 (H), H and L by 81.2 and 88.4 Å; for S19.8 (K), by 81.2 and 88.4 Å. MHC-I, β_2_m, and peptide COM in all complexes differed by less than 1.6 Å.

### AlphaFold-Multimer Prediction of Epitopes

While the crystallographic structure of each of the Fab/MHC complexes confirmed the general characteristics of the heavy/light chain associations of the Fv and C_L_C_H_ regions and of the fold and association of the MHC heavy chain with β_2_m, the final computational docking of the four Fabs to the pMHC/ β _2_m complexes contrasted starkly with the experimentally determined structures (Fig. 1, Fig. 5). While the Fab34-5-8 focuses on the α2 domain of H2-D^d^, as determined both from earlier exon-shuffling and mutagenic studies as well as the present X-ray structure, the AF-M predicted model docks this Fab to the α1 domain, focusing the V_H_V_L_ onto the opposite side of the molecule (Fig.5 A-C, gray).

The consistency of the X-ray structure of Fab34-2-12 bound to the α3 domain of H2-D^d^ is not borne out in the AF-M prediction (Fig. 5D-F). While the X-ray structure confirms the functionally defined reactivity (Fig. 2, 5D), the AF-M prediction placed the Fab interaction on the α1α2 domains, poised approximately like a TCR or TCR-like Ab (Fig. 5F). Once again, the final docking of a reasonably well-predicted Fab onto the well-predicted H-2D^d^ is at a completely different and non-overlapping site (Fig. 5E).

Fab28-14-8, a mAb that uniquely sees the α3 domain of H2-D^b^ and H2-L^d^, verified crystallographically (Fig. 3), was computationally docked to the TCR-like site involving the α1α2 and peptide surface (Fig. 5G-I). This docking placed Fab28-14-8 with its H chain poised over the α1 helix and its L chain over α2. This contrasts with the placement by AF-M of 34-2-12 with the Fab rotated about 90° to engage each chain with both helices (Fig. 5I). Again, the final docking performed by AF-M tended to focus the Fab roughly in a TCR or TCRm-like orientation (Fig. 5H). The final example of the test of the ability of AF-M to dock an Ab to its MHC/β_2_m antigen is that of FabS19.8. The X-ray structure of the complex (Fig. 4, Fig. 5J-L) clearly shows that the Fab, largely through contacts donated by its H chain, interacts predominantly with the β_2_m^b^(Ala85) MHC light chain, but also to six residues of the H2-D^d^ heavy chain. The Fab L chain sees some nine residues of both β_2_m and H2-D^d^. Remarkably, S19.8, although it binds a fully conformed H2-K^b^/β_2_m^b^ complex with a *K*_D_ of 1.3 × 10^-6^ M, also binds free β_2_m^b^ with an affinity approximately 6-fold weaker (Supplementary Fig. 1). AF-M placed the S19.8 Fab directly over the α1α2/peptide surface, again, much like a TCR (Fig. 5K-L), though experimentally, the Fab clearly focuses on β_2_m.

In summary, despite clear experimental (X-ray crystallographic) determination of the epitopic regions recognized by the four anti-MHC Fab studied here, AF-M in all cases failed to properly identify the antigenic surfaces identified by the antibodies. In three of the four cases, the AF-M dependent docking misconstrued the position of the Fab to be similar (but not identical) to that seen in dozens of TCR/MHC examples. Measurement of differences in the location of the center of mass of each of the chains in the superpositions is given in the legend to Fig. 5.

## DISCUSSION

The specificity of antibodies raised in precise genetic backgrounds, particularly in the mouse, has proven crucial in studies of antibody allotypes (*57*) and of polymorphism of genes controlling histocompatibility (*58*). The exploitation of mAb directed against human (*59*) and mouse (*28*) MHC antigens has revolutionized tissue typing for transplantation and our understanding of the genetic basis of immune responsiveness. Structural studies of the MHC class I (*60*) and class II molecules (*61, 62*) have contributed to a clear illustration of how MHC molecules bind peptide as a necessary prerequisite for presentation of antigenic peptides to CD4 and CD8 T cells. Exactly what regions of MHC molecules are bound by mAb that characterize particular domains or conformationally labile regions of MHC molecules remains in many cases poorly defined. Understanding with precision the nature of the epitopic sites seen by anti-MHC mAb may provide a basis for further engineering to optimize Ab/MHC. Here, we have characterized crystallographically the nature of four distinct regions of the mouse MHC-I molecules (Fig. 6A-B), including a peptide-dependent, but not peptide specific epitope, two related but distinct regions of the CD8-binding α3 domain, and a β_2_m allele-specific mAb. Careful inspection of the sites of interaction provides new insights into aspects of the peptide-dependent plasticity of the MHC-I molecule, the conserved site of α3 that interacts with CD8, or the location of a single residue polymorphism of β_2_m.

The structures of the four Fab/MHC-I complexes reported here reflect the domain organization of MHC-I molecules, peptide-dependent conformations of MHC-I, and MHC-I and β_2_m polymorphism. Early exon-shuffling experiments and subsequent mutagenesis and allele screening successfully identified domains of the molecules recognized—Fab34-5-8 binds the α2 domain, Fabs34-2-12 and 28-14-8 bind α3 although they approach the MHC molecule from rather different perspectives. The peptide dependency of 34-5-8 senses conformational changes in the 104-108 loop that result from peptide binding, even though this Fab itself does not interact with bound peptide. 28-14-8 addresses the α3 domain via a footprint that closely mimics that of the CD8αβ coreceptor ligand, while 34-2-12 approaches α3 via the membrane proximal loops, with a somewhat peripheral interaction with the CD8 binding loop of the α3 domain. S19.8, an anti-β_2_m mAb, is focused on a single amino acid polymorphism, and as expected, centers its antigen binding site on the polymorphic residue A85 of β_2_m. Although directed to A85 of β_2_m, S19.8 binds β_2_m both free and when in complex with an MHC-I heavy chain. Each of these four Fabs encounter residues that, on substitution, abolish tight interaction and thus account for their allelic specificity. Of the Fab/MHC-I complexes studied here, S19.8 serves as an example of a mAb that distinguishes a single residue substitution, A85 for D85 of β_2_m. The relatively low affinity of S19.8 for MHC-I/β_2_m (∼0.1 mM) suggests that the discrimination of the mAb is dependent on the lack of reactivity for the ASP85 variant rather than a strong association with Ala85. This result should also be considered in the context of a number of other antibodies that discriminate single residue differences, such as anti-allotype antibodies to immunoglobulins ((*63–65*)and antibodies that discriminate allelomorphs of cell surface receptors such as Thy-1 (*66*).

In addition to precise identification of the epitopes recognized by these four Fabs, the structures of the antibody/MHC complexes provide excellent test cases for the advanced prediction program AlphaFold2 and its computational progeny, AF-M. AlphaFold2 was highly successful in predicting the individual domains of the two murine peptide/MHC-I/β_2_m (H2-D^d^ and H2-D^b^) heterodimers studied and also of the four F_V_ segments of the Fab heterodimers examined. The interactions of the MHC-I with β_2_m predictions were considered acceptable to high (exhibiting DockQ scores greater than 0.8–see Supplementary Table 4), likely due to the large number of identical or similar experimental structural models in the protein database. The predictions of the interactions of the H and L chains of the Fab were acceptable, medium or high, and the docking scores of the four Fab binding to their pMHC antigens were incorrect (scores <0.2). Thus, when presented with sequences of heterodimers from a well-represented class of molecules, such as MHC-I, AlphaFold2-multimer performs well. Also, AF-M performed well with respect to the prediction of the basic fold of the Fv region of the four Fab, consistent with prior assessment of the veracity of such computational predictions (*67*). As expected, however, the backbone and side chains of the CDRH3 regions were less accurately ascertained (Supplementary Fig. 4). Most important, the ability of AF-M to predict the docking/binding of the Fab on the surface of the pMHC/β_2_m (i.e., the identification of the epitopic sites) was consistently incorrect.

Thus, our results examine several distinct aspects of the structure prediction algorithms, the ability: 1) to recognize and predict structure of individual domains; 2) to assemble and orient individual domains of a protein chain; 3) to generate the heterodimers of complexes frequently represented in the structural database; and finally 4) to produce models of the docking of the Fab V_H_/V_L_ heterodimer (paratope) onto its MHC-I heavy, heavy/β_2_m, or β_2_m epitope. Current views of antibody structure prediction recognize both the power of AlphaFold2 and its limitations, particularly in assessment of the CDR loops of antibodies (*67–69*). Protein antigen/antibody complexes continue to pose a major challenge for AI-powered prediction programs in several respects. Although the fold of the antibody itself is highly conserved and the framework of Fab structure is almost identical for all antibodies, the six antigen-binding sites established by the CDR loops that engage the epitope vary widely in length and composition. These six hypervariable loops are flexible and present a repertoire of dynamic states in solution that impose difficulties in accurately predicting a stabilized structure (*70*). In the cases we examined, although the folds of the Fv portion of the Fab were predicted correctly, the hinge angles between the variable and constant domains varied widely, and the CDR loops revealed a larger RMSD than the whole domains. The failure of AF-M to predict the epitopic sites (i.e. the detailed footprints of the Fab/MHC-I interactions) suggests that critical parameters such as binding energy or environmental conditions have been overlooked (*71*). Additional training with parameters such as elbow angles (*72*) or secondary structural features of epitopes (*73*) may help to improve the AI-based programs.

We must point out that predicting binding, epitopic, or docking sites in a complex structure is a process distinguished from predicting the structure of a single protein domain. The success of AlphaFold structural prediction depends on the advanced AI algorithm and high-performance computing. It also depends on the experimentally determined structures currently deposited in the PDB (∼200,000). AI training or deep learning from the PDB database in Alphafold2 creates ∼2 million parameters (*71*) in addition to the known structural parameters that greatly assist multiple sequence alignment for recognizing/constructing a similar structure from the PDB. AI training programs, such as Alphafold2, may evolve by learning additional parameters from the PDB as the number of experimental structures grows. Such training incorporates information from the experimental structures, but also may incur biases from related but distinct complex structures. For example, AF-M positioned three of the Fab in the Fab/MHC-I complexes (34-2-12, 28-14-8, and S19.8) to sit atop the α1α2 helices of the MHC-I, atop the peptide binding groove (Fig. 5F,5I,5L). There are many (more than 1000) TCR/MHC structures in the PDB (*74, 75*) in addition to several structures of complexes of TCRm/MHC (pdb 1W72(*76*), 3CVH(*77*), 4WUU(*78*),and 7TR4(*79*)). By and large, these structures indicate that the TCR or Fab binds over the α1α2 helices, while none of these complexes are similar to any of our mAb/MHC-I. Thus, AF-M predictions incurred a bias because of the available structures in the PDB. Once the Fab/MHC-I structures reported here are available in the PDB, they may be expected to contribute to improvement in the prediction of Fab/protein antigen complexes.

In summary, X-ray structures of mAb/protein antigens, as demonstrated here for a limited set of anti-MHC-I/MHC-I complexes, continue to provide detailed information describing the docking of Ab to their antigens, and explain the reactivity and specificity profiles of such Ab. As more mAb/Ag structures are experimentally determined, this should contribute to the elucidation of the parameters that determine mAb structure and Ag recognition, as we approach the goal of predicting antigenic specificity from Ab sequence.

## MATERIALS AND METHODS

### Source of mAb, production of Fab fragments, and production and purification of MHC-I molecules

Cells producing 34-5-8, 34-2-12, and 28-14-8 (originally designated 34-5-8S, 34-2-12S, and 28-14-8S, and occasionally referred to as 34.5.8, 34.2.12, and 28.14.8) were the gift of Drs. Keiko Ozato (NICHD, NIH) and David Sachs (NCI, NIH) (*28, 29*). 34-5-8 and 34-2-12 derive from BDF1 (H2^bxd^) anti-C3H (H2^k^) and 28-14-8 was derived from C3H.SW (H2^b^) anti-C3H/HeJ (H2^k^) responses. Cells producing S19.8 (SJL (H2^s^/β_2_m^a^) anti-B10.S (H2^s^/β_2_m^b^)) were obtained from Dr. Ulrich Hämmerling (Sloan-Kettering) (*30*). Cells were maintained in culture in Dulbecco’s modified Eagle’s medium, high glucose, supplemented with 10% fetal bovine serum, in 7.5% CO_2_ atmosphere at 37 °C. Cell culture supernatants were collected, purified by passage over Protein A Sepharose (Cytiva) washed with 0.45 M NaCl, 10 mM Tris pH 8.0, eluted with 0.1 M glycine HCl pH 3.0, into 1 M Tris pH 8.0, and dialyzed against 1X PBS. Fab fragments were then prepared and purified following papain digestion, protein A purification to remove undigested molecules and Fc fragments, followed by size exclusion chromatography on either Superdex 200 or Superdex 200 increase columns (Cytiva). *In vitro* expressed and refolded H2-D^d^,-D^b^ were prepared with β_2_m^b^ and peptides RGPGRAFVTI (HIVIIIB glycoprotein peptide, for H2-D^d^) and ASNENMETM (influenza peptide, for H2-D^b^) as described previously (*80*) .

### Preparation of mRNA, cDNA, and sequence determination of mAb encoding H and L chains

Total RNA was extracted from 10^7^ hybridoma cells using the Monarch^TM^ total RNA extraction kit (New England BioLabs, Ipswich, MA, USA) following manufacturer’s instructions. Two μg of RNA served as template for cDNA synthesis using oligo dT and murine leukemia reverse transcriptase as implemented in the OneTaq RT PCR kit (New England Biolabs). A panel of oligonucleotides designed to amplify mouse Ig V genes as described by Wang et al (*81*) was used to PCR amplify and then sequence V_H_ and V_L_ from each hybridoma. The encoded protein sequences are shown in Supplementary Figure 5.

### Preparation of complexes, X-ray crystallization conditions, data collection, and refinement

Equimolar amounts of purified Fab and MHC proteins were incubated at 25 °C for 2-3 hours. Complexes were isolated on a Superdex 200 increase column in 1X PBS, concentrated and buffer exchanged into 25 mM Tris, pH 8.0, 50 mM NaCl in preparation for crystallization.

Crystallization conditions were identified by screening hanging drops at 18 °C. Crystals of Fab28-14-8/H2-D^b^ were grown in 16% PEG 4000, 0.1 M HEPES pH 7.5, 0.2 M MgCl. Crystals of Fab34-5-8/H2-D^d^ were grown in 18% PEG 4000, 0.1 M MES, pH 6.0, 0.12 M Ca Acetate. Crystals of Fab34-2-12/H2-D^d^ were obtained in 12% PEG 8000, 0.1 M MES, pH 6.5. Crystals of FabS19.8/H2-D^d^ grew in 0.5 M Ammonium Sulfate, 0.1 M Na citrate, pH 5.6, 1.0 M LiSO_4_ and were further improved by seeding. Crystals were cryoprotected in mother liquor containing 10% ethylene glycol, and flash frozen in liquid nitrogen. Diffraction data were collected (at wavelength 1.033 Å, in N_2_ stream at ∼100 K) at Southeast Regional Collaborative Access Team (SER-CAT) beamline 22ID at the Advanced Photon Source, Argonne National Laboratory and processed with XDS (*82*) to 2.6 Å, 2.7 Å, 2.8 Å and 2.9 Å resolution for Fab28-14-8/H2-D^b^, Fab34-5-8/H2-D^d^, Fab34-2-12/H2-D^d^, and FabS19.8/H2-D^d^/β_2_m^b^ respectively (see Table 1). The structures were solved by molecular replacement with Phaser (*83*) using H2-D^b^ from PDB 1WBX or H2-D^d^ from PDB 5WEU as search models. For the Fab search we used a model of DX17 Fab (not yet deposited) with the CDR loops trimmed as the initial search model. Molecular replacement solutions were subjected to several rounds of refinement with Phenix (*84, 85*) interspersed with manual building in Coot (*86*). R_work_/R_free_ (%) values for final refined models of Fab28-14-8/H2-D^b^, Fab34-5-8/H2-D^d^, Fab34-2-12/H2-D^d^, and FabS19.8/H2-D^d^ are 21.5/25.5, 22.7/25.7, 23.5/28.6, and 19.2/24.9 respectively. Data collection and refinement statistics are summarized in Table 1. Graphics figures were generated with PyMOL (*87*) and ChimeraX (*38, 39*).

### Structure analysis and computational predictions

Analysis of X-ray structures and comparison of computational predictions with experimental structures was carried out with a variety of programs, including *S_c_* (*45*), CNS1.3 (*88*), PDBePISA (*89*), PyMOL(*87*), and ChimeraX(*39*). Antibody V_H_ and V_L_ sequences, determined from cDNA as described above, and the corresponding MHC-I and β_2_m sequences were entered into the AlphaFold structure prediction module of ChimeraX 1.6 (Tools>Structure Prediction>AlphaFold>Predict) which queried Colab via Google servers. The resulting “best_model” was further analyzed and compared with our X-ray structure of the same complex. In cases where the X-ray structure contained more than one complex in the asymmetric unit (ASU), the first complex was used. Center of mass (COM) for indicated chains was calculated in ChimeraX (*39*).

### Surface plasmon resonance (SPR)

SPR experiments were performed as described previously (*90*) at 25 °C on a Biacore^TM^ T200 (Cytiva, Uppsala, Sweden) in 10 mM HEPES pH 7.4, 150 mM NaCl, 3 mM EDTA, and 0.05% Tween-20 at a flow rate of 30 ml/min. mAb S19.8 (Santa Cruz Technology, Catalog #SC-32241) was repurified on a Superdex 200 increase column (Cytiva) in PBS, and 1100 resonance units (RU) were immobilized on a series S CM5 sensor chip (Cytiva) by amin (NHS-EDC) coupling. A reference cell was mock coupled to allow for background subtraction. Various MHC-I proteins described in figure legends were prepared with either human β_2_m or mouse β_2_m^b^. The binding surface was regenerated with 5 mM phosphoric acid. Kinetics studies were performed with graded concentrations as indicated in the figure legends. Sensorgrams were globally fitted to a 1:1 binding model as implemented in Biacore^TM^ T200 Evaluation Software v3.1 and plotted with GraphPad Prism. Proteins used in SPR were HLA-A*02:01/h β_2_m/flu peptide, HLA-A*02:01/mβ_2_m^b^/flu peptide, H2-K^b^/hβ_2_m/ova peptide, H2-K^b^/mβ_2_m^b^/ova peptide, mβ_2_m^b^ or human β_2_m. Flu peptide is GILGFVFTL, representing influenza matrix protein M1 (58–66) and ova peptide is SIINFEKL, ovalbumin residues 257-264. The indicated complexes were refolded from bacterially expressed inclusion bodies and synthetic peptide and purified by standard methods (*80*).

## ACKNOWLEDGEMENTS

This work was supported by the Intramural Research Program of the NIAID, NIH, projects **ZIA AI000394-40** and **ZIA AI000622-32.** We appreciate the comments of John Altman, Christopher Boughter, Ted Hansen, Eduardo Padlan, T.V. Rajan, and Sebastian Springer. X-ray data were collected at SER-CAT (22-ID) and GM/CA-CAT (23ID) beamlines at the Advanced Photon Source (APS), Argonne National Laboratory. Use of the APS was supported by the U.S. Department of Energy, Office of Science, Office of Basic Energy Sciences, under Contract No. W-31-109-Eng-38. GM/CA@APS has been funded in whole or in part with Federal funds from the National Cancer Institute (ACB-12002) and the National Institute of General Medical Sciences (AGM-12006). Author contributions: Conceptualization: LFB, JJ, KN, DHM; Methodology: JJ, DHM; Investigation: LFB, JJ, JA, KN, DHM; Computational analysis: JJ, JA, KN; Visualization: LFB, JJ, DHM; Funding Acquisition: DHM; Project Administration: DHM; Supervision: DHM; Writing – original draft: DHM; Scripting – review & editing: LFB, JJ, JA, KN, DHM. Authors declare that they have no competing interests. Model coordinates and structure factors are deposited with the RCSB under PDB accession numbers 8TQA, 8, 8, and 8 for Fab28-14-8, Fab34-5-8, Fab34-2-12, and FabS19.8 respectively. Plasmid vectors may be obtained from the authors under NIH MTA.

## REFERENCES AND NOTES

1. G. Kohler, C. Milstein, Continuous cultures of fused cells secreting antibody of predefined specificity. Nature 256, 495–497 (1975).

2. B. A. Diamond, D. E. Yelton, M. D. Scharff, Monoclonal antibodies. A new technique for producing serologic reagents. N Engl J Med 304, 1344–1349 (1981).

3. M. D. Scharff, S. Roberts, P. Thammana, Monoclonal antibodies. J Infect Dis 143, 346–351 (1981).

4. G. J. Hammerling et al., Monoclonal antibodies against murine cell surface antigens: anti-H-2, anti-Ia and anti-T cell antibodies. Curr Top Microbiol Immunol 81, 100–106 (1978).

5. H. Lemke, G. J. Hammerling, U. Hammerling, Fine specificity analysis with monoclonal antibodies of antigens controlled by the major histocompatibility complex and by the Qa/TL region in mice. Immunol Rev 47, 175–206 (1979).

6. V. T. Oi, P. P. Jones, J. W. Goding, L. A. Herzenberg, L. A. Herzenberg, Properties of monoclonal antibodies to mouse Ig allotypes, H-2, and Ia antigens. Curr Top Microbiol Immunol 81, 115–120 (1978).

7. F. M. Brodsky, P. Parham, C. J. Barnstable, M. J. Crumpton, W. F. Bodmer, Monoclonal antibodies for analysis of the HLA system. Immunol Rev 47, 3–61 (1979).

8. K. C. Stallcup, T. A. Springer, M. F. Mescher, Characterization of an anti-H-2 monoclonal antibody and its use in large-scale antigen purification. J Immunol 127, 923–930 (1981).

9. S. H. Herrmann, M. F. Mescher, Purification of the H-2Kk molecule of the murine major histocompatibility complex. J Biol Chem 254, 8713–8716 (1979).

10. J. Colombani, V. Lepage, C. Raffoux, M. Colombani, HLA typing with monoclonal antibodies: evaluation of 356 HLA monoclonal antibodies including 181 studied during the 10th International Histocompatibility Workshop. Tissue antigens 34, 97–110 (1989).

11. J. S. Blum, P. A. Wearsch, P. Cresswell, Pathways of antigen processing. Annu Rev Immunol 31, 443–473 (2013).

12. K. L. Rock, E. Reits, J. Neefjes, Present Yourself! By MHC Class I and MHC Class II Molecules. Trends Immunol 37, 724–737 (2016).

13. M. I. J. Raybould, D. A. Nissley, S. Kumar, C. M. Deane, Computationally profiling peptide:MHC recognition by T-cell receptors and T-cell receptor-mimetic antibodies. Front Immunol 13, 1080596 (2022).

14. R. Frick, et al., A high-affinity human TCR-like antibody detects celiac disease gluten peptide-MHC complexes and inhibits T cell activation. Sci Immunol 6, (2021).

15. K. Ozato, S. L. Epstein, P. Henkart, T. H. Hansen, D. H. Sachs, Studies on monoclonal antibodies to mouse MHC products. Transplant Proc 13, 958–962 (1981).

16. B. D. Tait, Detection of HLA Antibodies in Organ Transplant Recipients - Triumphs and Challenges of the Solid Phase Bead Assay. Front Immunol 7, 570 (2016).

17. G. A. Evans, D. H. Margulies, B. Shykind, J. G. Seidman, K. Ozato, Exon shuffling: mapping polymorphic determinants on hybrid mouse transplantation antigens. Nature 300, 755–757 (1982).

18. D. H. Margulies, J. McCluskey, Exon shuffling: new genes from old. Surv Immunol Res 4, 146–159 (1985).

19. V. H. Engelhard, J. R. Yannelli, G. A. Evans, S. F. Walk, M. J. Holterman, Construction of novel class I histocompatibility antigens by interspecies exon shuffling. J Immunol 134, 4218–4225 (1985).

20. H. Allen, D. Wraith, P. Pala, B. Askonas, R. A. Flavell, Domain interactions of H-2 class I antigens alter cytotoxic T-cell recognition sites. Nature 309, 279–281 (1984).

21. T. A. Potter, J. A. Bluestone, T. V. Rajan, A single amino acid substitution in the alpha 3 domain of an H-2 class I molecule abrogates reactivity with CTL. J Exp Med 166, 956–966 (1987).

22. D. H. Mattson et al., Differential effects of amino acid substitutions in the beta-sheet floor and alpha-2 helix of HLA-A2 on recognition by alloreactive viral peptide-specific cytotoxic T lymphocytes. J Immunol 143, 1101–1107 (1989).

23. D. H. Hausmann, B. Yu, S. Hausmann, K. W. Wucherpfennig, pH-dependent peptide binding properties of the type I diabetes-associated I-Ag7 molecule: rapid release of CLIP at an endosomal pH. J Exp Med 189, 1723–1734 (1999).

24. M. Waldenstrom, A. Achour, J. Michaelsson, A. Rolle, K. Karre, The role of an exposed loop in the alpha(2) domain in the mouse MHC class IH-2D(d) molecule for recognition by the monoclonal antibody 34-5-8S and the NK-cell receptor Ly49A. Scand J Immunol 55, 129–139 (2002).

25. K. Ozato, G. A. Evans, B. Shykind, D. H. Margulies, J. G. Seidman, Hybrid H-2 histocompatibility gene products assign domains recognized by alloreactive T cells. Proc Natl Acad Sci U S A 80, 2040–2043 (1983).

26. F. M. Karlhofer, R. K. Ribaudo, W. M. Yokoyama, MHC class I alloantigen specificity of Ly-49+ IL-2-activated natural killer cells. Nature 358, 66–70 (1992).

27. A. K. Panda et al., Cutting Edge: Inhibition of the Interaction of NK Inhibitory Receptors with MHC Class I Augments Antiviral and Antitumor Immunity. J Immunol 205, 567–572 (2020).

28. K. Ozato, N. M. Mayer, D. H. Sachs, Monoclonal antibodies to mouse major histocompatibility complex antigens. Transplantation 34, 113–120 (1982).

29. K. Ozato, T. H. Hansen, D. H. Sachs, Monoclonal antibodies to mouse MHC antigens. II. Antibodies to the H-2Ld antigen, the products of a third polymorphic locus of the mouse major histocompatibility complex. J Immunol 125, 2473–2477 (1980).

30. N. Tada, S. Kimura, A. Hatzfeld, U. Hammerling, Ly-m11: the H-3 region of mouse chromosome 2 controls a new surface alloantigen. Immunogenetics 11, 441–449 (1980).

31. D. H. Margulies, J. R. Parnes, N. A. Johnson, J. G. Seidman, Linkage of beta 2-microglobulin and ly-m11 by molecular cloning and DNA-mediated gene transfer. Proc Natl Acad Sci U S A 80, 2328–2331 (1983).

32. R. Evans et al., Protein complex prediction with AlphaFold-Multimer. bioRXiv, (2022).

33. J. Jumper et al., Highly accurate protein structure prediction with AlphaFold. Nature 596, 583–589 (2021).

34. A. O. Stevens, Y. He, Benchmarking the Accuracy of AlphaFold 2 in Loop Structure Prediction. Biomolecules 12, (2022).

35. A. David, S. Islam, E. Tankhilevich, M. J. E. Sternberg, The AlphaFold Database of Protein Structures: A Biologist’s Guide. J Mol Biol 434, 167336 (2022).

36. R. Yin, B. G. Pierce, Evaluation of AlphaFold Antibody-Antigen Modeling with Implications for Improving Predictive Accuracy. bioRxiv, (2023).

37. F. Wong et al., Benchmarking AlphaFold-enabled molecular docking predictions for antibiotic discovery. Mol Syst Biol 18, e11081 (2022).

38. T. D. Goddard et al., UCSF ChimeraX: Meeting modern challenges in visualization and analysis. Protein Sci 27, 14–25 (2018).

39. E. F. Pettersen et al., UCSF ChimeraX: Structure visualization for researchers, educators, and developers. Protein Sci 30, 70–82 (2021).

40. M. Mirdita et al., ColabFold: making protein folding accessible to all. Nat Methods 19, 679–682 (2022).

41. G. R. Otten et al., Peptide and beta 2-microglobulin regulation of cell surface MHC class I conformation and expression. J Immunol 148, 3723–3732 (1992).

42. J. P. Abastado et al., Fine mapping of epitopes by intradomain Kd/Dd recombinants. J Exp Med 166, 327–340 (1987).

43. C. Murre et al., Construction, expression and recognition of an H-2 molecule lacking its carboxyl terminus. Nature 307, 432–436 (1984).

44. J. Sundback et al., The alpha2 domain of H-2Dd restricts the allelic specificity of the murine NK cell inhibitory receptor Ly-49A. J Immunol 160, 5971–5978 (1998).

45. W. P. Coleman, 3rd, N. Lawrence, R. N. Sherman, R. J. Reed, K. S. Pinski, Autologous collagen? Lipocytic dermal augmentation. A histopathologic study. J Dermatol Surg Oncol 19, 1032–1040 (1993).

46. N. Matsumoto, W. M. Yokoyama, S. Kojima, K. Yamamoto, The NK cell MHC class I receptor Ly49A detects mutations on H-2Dd inside and outside of the peptide binding groove. J Immunol 166, 4422–4428 (2001).

47. R. J. Rubocki et al., Mutation at amino acid position 133 of H-2Dd prevents beta 2m association and immune recognition but not surface expression. J Immunol 146, 2352–2357 (1991).

48. R. Wang, K. Natarajan, D. H. Margulies, Structural basis of the CD8 alpha beta/MHC class I interaction: focused recognition orients CD8 beta to a T cell proximal position. J Immunol 183, 2554–2564 (2009).

49. A. K. Mitra et al., Supine orientation of a murine MHC class I molecule on the membrane bilayer. Curr Biol 14, 718–724 (2004).

50. J. McCluskey, J. A. Bluestone, J. E. Coligan, W. L. Maloy, D. H. Margulies, Serologic and T cell recognition of truncated transplantation antigens encoded by in vitro deleted class I major histocompatibility genes. J Immunol 136, 1472–1481 (1986).

51. J. McCluskey, R. N. Germain, D. H. Margulies, Cell surface expression of an in vitro recombinant class II/class I major histocompatibility complex gene product. Cell 40, 247–257 (1985).

52. J. M. Connolly, T. H. Hansen, A. L. Ingold, T. A. Potter, Recognition by CD8 on cytotoxic T lymphocytes is ablated by several substitutions in the class I alpha 3 domain: CD8 and the T-cell receptor recognize the same class I molecule. Proc Natl Acad Sci U S A 87, 2137–2141 (1990).

53. H. Allen, J. Fraser, D. Flyer, S. Calvin, R. Flavell, Beta 2-microglobulin is not required for cell surface expression of the murine class I histocompatibility antigen H-2Db or of a truncated H-2Db. Proc Natl Acad Sci U S A 83, 7447–7451 (1986).

54. E. Hermel, P. J. Robinson, J. X. She, K. F. Lindahl, Sequence divergence of B2m alleles of wild Mus musculus and Mus spretus implies positive selection. Immunogenetics 38, 106–116 (1993).

55. P. J. Robinson, M. Steinmetz, K. Moriwaki, K. F. Lindahl, Beta-2 microglobulin types in mice of wild origin. Immunogenetics 20, 655–665 (1984).

56. S. Basu, B. Wallner, DockQ: A Quality Measure for Protein-Protein Docking Models. PLoS One 11, e0161879 (2016).

57. R. Lieberman, S. Rudikoff, W. Humphrey, Jr., M. Potter, Allelic forms of anti-phosphorylcholine antibodies. J Immunol 126, 172–176 (1981).

58. G. D. Snell, Recent advances in histocompatibility immunogenetics. Adv Genet 20, 291–355 (1979).

59. P. Parham, W. F. Bodmer, Monoclonal antibody to a human histocompatibility alloantigen, HLA-A2. Nature 276, 397–399 (1978).

60. P. J. Bjorkman, P. Parham, Structure, function, and diversity of class I major histocompatibility complex molecules. Annu Rev Biochem 59, 253–288 (1990).

61. J. H. Brown et al., Three-dimensional structure of the human class II histocompatibility antigen HLA-DR1. Nature 364, 33–39 (1993).

62. L. J. Stern et al., Crystal structure of the human class II MHC protein HLA-DR1 complexed with an influenza virus peptide. Nature 368, 215–221 (1994).

63. R. Grubb, The Gm system. Anti-Gm’s: characteristics in rheumatoid arthritis; experimental induction without resort to allotype; frequent occurrence in mononucleosis. Scand J Rheumatol Suppl 75, 227–232 (1988).

64. N. McCartney-Francis, R. M. Skurla, Jr., R. G. Mage, K. E. Bernstein, Kappa-chain allotypes and isotypes in the rabbit: cDNA sequences of clones encoding b9 suggest an evolutionary pathway and possible role of the interdomain disulfide bond in quantitative allotype expression. Proc Natl Acad Sci U S A 81, 1794–1798 (1984).

65. R. Riblet et al., Genetics of mouse antibodies. I. Linkage of the dextran response locus, VH-DEX, to allotype. Eur J Immunol 5, 775–777 (1975).

66. A. F. Williams, J. Gagnon, Neuronal cell Thy-1 glycoprotein: homology with immunoglobulin. Science 216, 696–703 (1982).

67. M. L. Fernandez-Quintero et al., Challenges in antibody structure prediction. MAbs 15, 2175319 (2023).

68. B. Abanades et al., ImmuneBuilder: Deep-Learning models for predicting the structures of immune proteins. Commun Biol 6, 575 (2023).

69. M. L. Fernandez-Quintero et al., CDR loop interactions can determine heavy and light chain pairing preferences in bispecific antibodies. MAbs 14, 2024118 (2022).

70. M. L. Fernandez-Quintero et al., Paratope states in solution improve structure prediction and docking. Structure 30, 430–440 e433 (2022).

71. T. C. Terwilliger et al., Improved AlphaFold modeling with implicit experimental information. Nat Methods 19, 1376–1382 (2022).

72. R. L. Stanfield, A. Zemla, I. A. Wilson, B. Rupp, Antibody elbow angles are influenced by their light chain class. J Mol Biol 357, 1566–1574 (2006).

73. J. Jiang et al., SARS-CoV-2 antibodies recognize 23 distinct epitopic sites on the receptor binding domain. Commun Biol 6, 953 (2023).

74. P. Marrack, J. P. Scott-Browne, S. Dai, L. Gapin, J. W. Kappler, Evolutionarily conserved amino acids that control TCR-MHC interaction. Annu Rev Immunol 26, 171–203 (2008).

75. B. M. Baker, D. R. Scott, S. J. Blevins, W. F. Hawse, Structural and dynamic control of T-cell receptor specificity, cross-reactivity, and binding mechanism. Immunol Rev 250, 10–31 (2012).

76. M. Hulsmeyer et al., A major histocompatibility complex-peptide-restricted antibody and t cell receptor molecules recognize their target by distinct binding modes: crystal structure of human leukocyte antigen (HLA)-A1-MAGE-A1 in complex with FAB-HYB3. J Biol Chem 280, 2972–2980 (2005).

77. T. Mareeva, E. Martinez-Hackert, Y. Sykulev, How a T cell receptor-like antibody recognizes major histocompatibility complex-bound peptide. J Biol Chem 283, 29053–29059 (2008).

78. N. Ataie et al., Structure of a TCR-Mimic Antibody with Target Predicts Pharmacogenetics. J Mol Biol 428, 194–205 (2016).

79. X. Yang et al., Facile repurposing of peptide-MHC-restricted antibodies for cancer immunotherapy. Nat Biotechnol 41, 932–943 (2023).

80. H. Li, K. Natarajan, E. L. Malchiodi, D. H. Margulies, R. A. Mariuzza, Three-dimensional structure of H-2Dd complexed with an immunodominant peptide from human immunodeficiency virus envelope glycoprotein 120. J Mol Biol 283, 179–191 (1998).

81. Z. Wang et al., Universal PCR amplification of mouse immunoglobulin gene variable regions: the design of degenerate primers and an assessment of the effect of DNA polymerase 3’ to 5’ exonuclease activity. J Immunol Methods 233, 167–177 (2000).

82. W. Kabsch, Xds. Acta Crystallogr D Biol Crystallogr 66, 125–132 (2010).

83. A. J. McCoy et al., Phaser crystallographic software. J Appl Crystallogr 40, 658–674 (2007).

84. P. D. Adams et al., PHENIX: a comprehensive Python-based system for macromolecular structure solution. Acta Crystallogr D Biol Crystallogr 66, 213–221 (2010).

85. D. Liebschner et al., Macromolecular structure determination using X-rays, neutrons and electrons: recent developments in Phenix. Acta Crystallogr D Struct Biol 75, 861–877 (2019).

86. P. Emsley, B. Lohkamp, W. G. Scott, K. Cowtan, Features and development of Coot. Acta Crystallogr D Biol Crystallogr 66, 486–501 (2010).

87. PyMOL.

88. A. T. Brunger et al., Crystallography & NMR system: A new software suite for macromolecular structure determination. Acta Crystallogr D Biol Crystallogr 54, 905–921 (1998).

89. E. Krissinel, K. Henrick, Inference of macromolecular assemblies from crystalline state. J Mol Biol 372, 774–797 (2007).

90. J. Jiang et al., Structural mechanism of tapasin-mediated MHC-I peptide loading in antigen presentation. Nat Commun 13, 5470 (2022).

91. F. Madeira et al., Search and sequence analysis tools services from EMBL-EBI in 2022. Nucleic Acids Res 50, W276–W279 (2022).

92. X. Robert, P. Gouet, Deciphering key features in protein structures with the new ENDscript server. Nucleic Acids Res 42, W320–324 (2014).

