## Supplementary Materials for "Experimental structures of antibody/MHC-I complexes reveal details of epitopes overlooked by computational prediction"

SUPPLEMENTARY FIGURE 1.

SUPPLEMENTARY FIGURE 2.

SUPPLEMENTARY FIGURE 3.

SUPPLEMENTARY FIGURE 4.

SUPPLEMENTARY FIGURE 5.

SUPPLEMENTARY TABLE 1.

SUPPLEMENTARY TABLE 2.

SUPPLEMENTARY TABLE 3.

SUPPLEMENTARY TABLE 4.

Supplementary Fig. 1

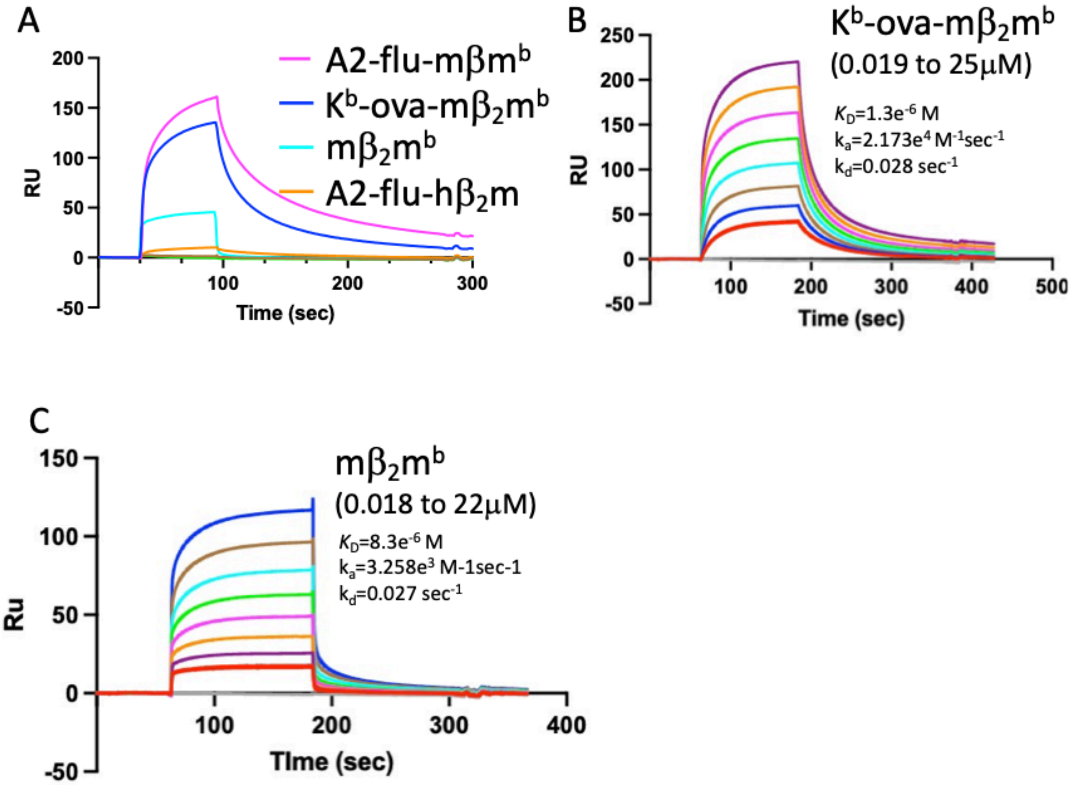

Supplementary Figure 1. S19.8 binds MHC-I molecules complexed with mouse $\beta_2 m^b$ but not with human $\beta_2$m. (A) SPR analysis was performed as described in Methods using a surface coupled with S19.8 (1100 RU), using preparations of pMHC consisting of: HLA-A*02:01/flu peptide/human $\beta_2$m or mouse $\beta_2 m^b$, H2-$K^b$/ova peptide (SIINFEKL)/human $\beta_2$m or mouse $\beta_2 m^b$, or free human $\beta_2$m or mouse $\beta_2 m^b$ at 1 uM. Baseline tracings reveal the lack of binding of free human $\beta_2$m (green) or of H2-$K^b$-ova-h$\beta_2$m. (B) Graded concentrations (from 0.019 to 25 uM) of H2-$K^b$-SIINFEKL-$\beta_2 m^b$ were offered to S19.8. (C) Graded concentrations (from 0.18 to 22 uM) of mouse $\beta_2 m^b$ were offered to the S19.8 surface. Curves were fit globally to a single site kinetics model using BiaEval 3.0. Indicated parameters are representative of three independent experiments with different preparations of S19.8.

### Supplementary Fig.2

### A

| Complex | X-ray model | AF-M model | MHC-I (A+B+P) | A (α-chain) | B ($\beta_2$m) | P (pep) | Loop | MHC-I α1α2 | MHC-I α3 |
|---|---|---|---|---|---|---|---|---|---|
| 34-5-8/$D^d$ | 8TQ8 | 8TQ8-AF | 1.814 | 1.330 | 2.690 | 2.574 | --- | 1.259 | 1.456 |
| 34-2-12/$D^d$ | 8TQ7 | 8TQ7-AF | 1.360 | 1.369 | 1.278 | 2.101 | --- | 1.272 | 1.534 |
| 28-14-8/$D^b$ | 8TQA | 8TQA-AF | 1.793 | 1.931 | 1.331 | 1.859 | 5.428 | 2.099 | 1.558 |
| S19.8/$D^d$/ $\beta_2m^b$ | 8TQ9 | 8TQ9-AF | 1.703 | 1.732 | 1.422 | 3.168 | 5.164 | 1.664 | 1.855 |

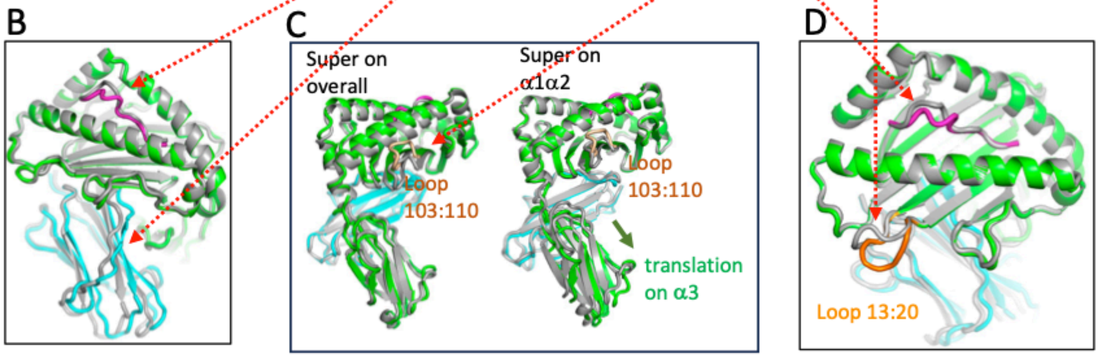

Supplementary Figure 2. Comparison of X-ray and AF-M models of pMHC in different complexes. The indicated Fab/MHC-I complexes were predicted by AF-M as described in the Methods and compared with the indicated refined crystallographic models. (A) Comparisons (reported as Å) were performed as all-atom RMSD using CNS 1.3 as described in Materials and Methods. All three chains (A+B+P), each protein chain alone (α, H chain; $\beta_2$m L chain), or individual domain units were compared as indicated. The hinge angle defined by α1α2 vs α3, was also calculated. (B) Superposition of H2-$D^d$ from 8TQ8 as compared with the pMHC from the AF-M best_model for H2-$D^d$ from the Fab34-5-8 complex. (C) Superposition of H2-$D^b$ from 8TQA as compared with the pMHC from the AF-M best_model for H2-$D^b$ from the Fab28-14-8 complex;

and (D) Superposition of H2-$D^d$ from 8TQ9 as compared with the pMHC from the AF-M best_model for H2-$D^d$ from the FabS19.8 complex.

Supplementary Fig. 3.

### RMSD vs residue no. – X-ray and AF-M models

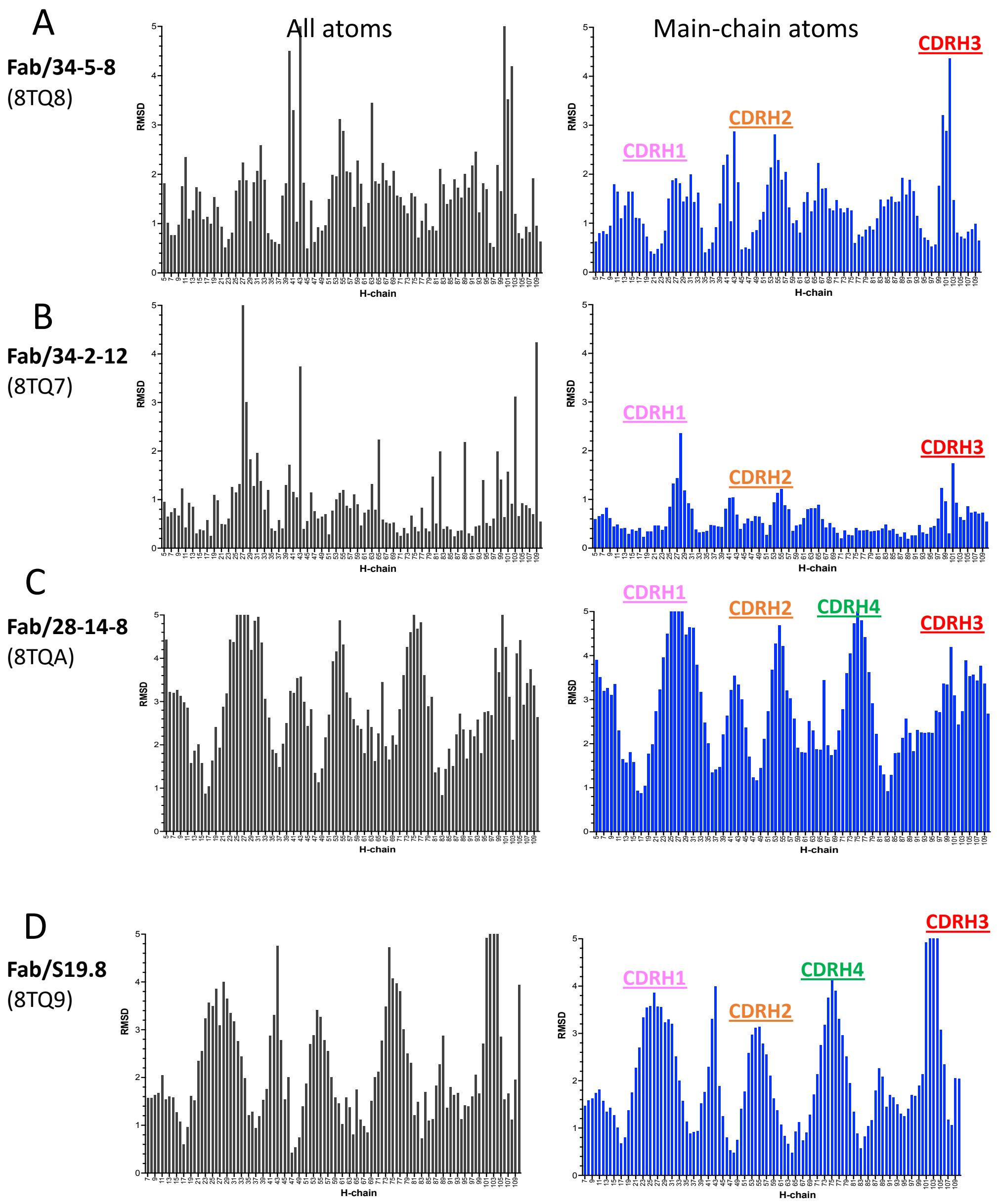

Supplementary Figure 3. Comparison of RMSD vs residue number reveals major differences in CDR loops. RMSD values determined for all atoms (left) and main-chain atoms (right) for the indicated four mAb/MHC complexes as determined crystallographically vs by AF-M.

### Supplementary Figure 4.

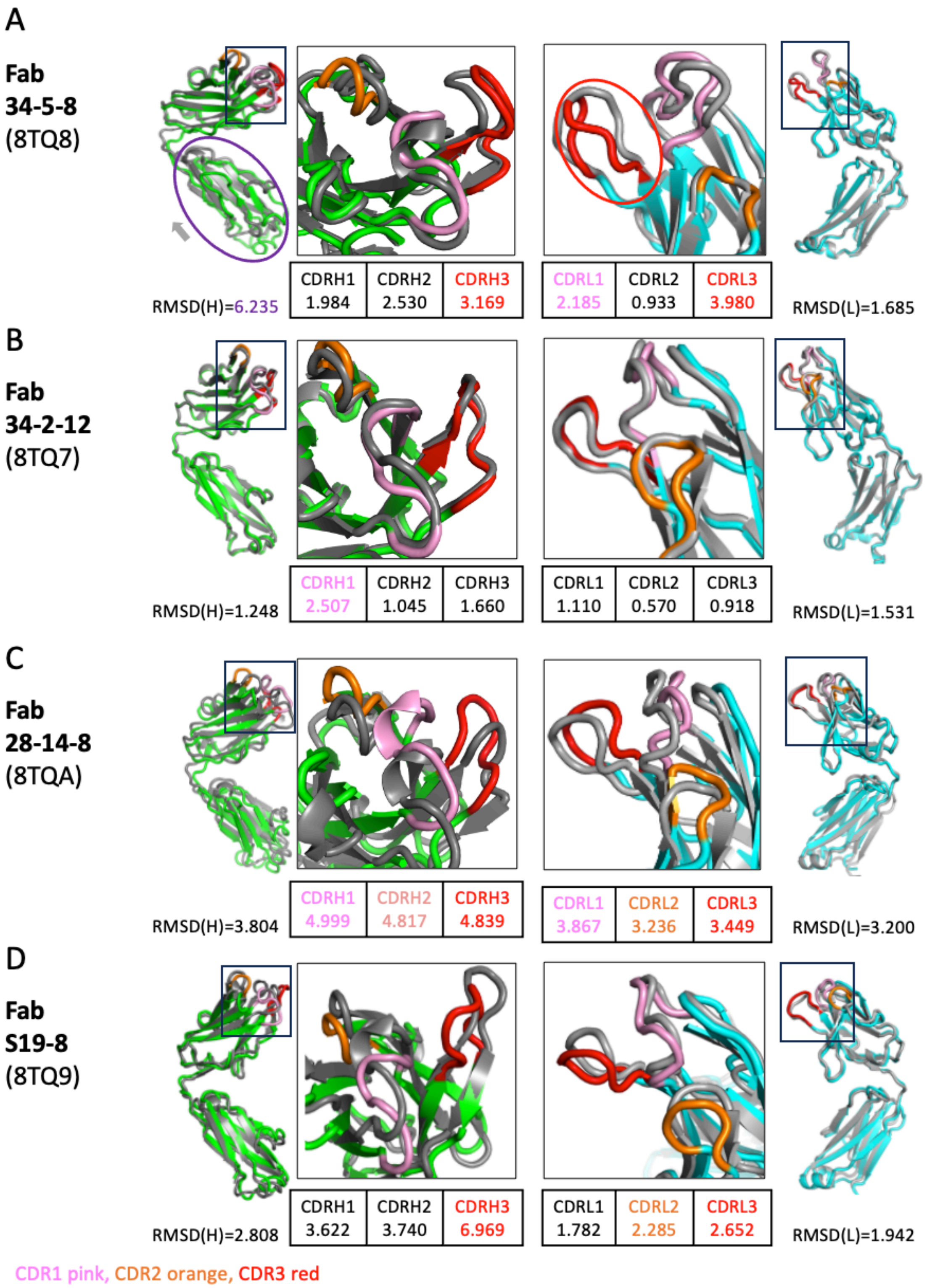

Supplementary Figure 4. Illustration of major differences in conformation of CDR loops of H chains (left) and L chain (right). H or L chain comparing X-ray to AF-M structures. Superpositions of H or L chains were accomplished in PyMOL and RMSD of CDR loops were calculated as shown in illustrations or the accompanying tables.

### Supplementary Fig. 5.

**A**

**H-Chain**

```
Fab3458-H       ------EESGPELGKPGASVKMSCTASGYAFTSYVMHWVMQKPGQGLEWIGYFNPYNDAG  58
Fab34212-H      ---VQLQQSGPELVKPGASVKISCQSSGYTISNSWMNWVKQRPGKGLEWIGRIYPGDGDT  58
Fab2814S-H      ------QQSGAELAKPGASVKMSCKASGYNFTTYWMHWVRQRPGQGLEWIGYINPSSGYT  58
FabS19-H        ---VQLQQSGPVLVKPGASVKMSCKASGYTFTDHYMNWVKQSPGKSLEWIGGINPYNGGI  58

Fab3458-H       KYNAKFKGKATLTSDKSSNTAYMELSSLTSEDSTVYYCARA--Y-FSKGPFAYWGQGTLV  115
Fab34212-H      HYNGKFKAKATLTADRSSSTAYMHLSSLTSEDSAIYFCARSTA-----AWFPYWGRGTLV  113
Fab2814S-H      EFNQKFKGKATLTADKSSNTAYIQLTSLTSDDSAIYYCARSF------YYFDYWGQGTTL  112
FabS19-H        SYNQKFKGMATLTADKSSSTAYMELNSLTSEDSAVYYCVRRIGY-YPSYFFDYWGQGTTL  116
```

**B**

**L-Chain**

```
Fab3458-L       ----TQTPASLAVSLGQRATISCRASESVDRHGNSFMHWYQQKPGQPPKLLIYRASNLDS  60
Fab34212-L      -TTVTQTPKFMSTSVGDRVSVTCTASQNV----DTNVAWYQQKPGQSPKPLIFSASSRYT  56
Fab2814S-L      ----TQTPSSLSASLGDRVTISCRASQDI----SNYLNWYQQKPDGPVKLLIYYTSRLHS  56
FabS19-L        -TTVTQSPASLSVATGEKVTIRCITSTDI----DDNMNWYQQKPGEPPKLLISEGNTLRP  56

Fab3458-L       GIPARFSGSGSRTDFTLTINPVEADDVATYYCQQSNEDPPWTFGGGTKLEIKRADAAPTV  120
Fab34212-L      GIPDRFGGSGSGTDFALTISNVQSEDLAEYFCQQYH-SFPYTFGGGTRLEIKRADAAPTV  115
Fab2814S-L      GVPSRFSGSGSGTDYSLTISNLEQEDIATYFCQQGN-TFPWTFGGGTKLEIKRADAAPTV  115
FabS19-L        GVPSRFSSSGFGTDFVFTIENTLSEDFADYYCLQSD-DMPLTFGAGTKLELKRADAAPTV  115
```

**C**

8TQ8

| CDR | 34-5-8 |
|---|---|
| CDRH1 | S25-V33 |
| CDRH2 | N52-D56 |
| CDRH3 | R98-Y108 |
| CDRL1 | S26-F36 |
| CDRL2 | R54-L58 |
| CDRL3 | Q94-W101 |

8TQ7

| CDR | 34-2-12 |
|---|---|
| CDRH1 | S25-M33 |
| CDRH2 | Y52-D57 |
| CDRH3 | R98-Y106 |
| CDRL1 | S26-V33 |
| CDRL2 | F49-S53 |
| CDRL3 | Q90-Y96 |

8TQA

| CDR | 28-14-8 |
|---|---|
| CDRH1 | S25-W33 |
| CDRH2 | N52-G56 |
| CDRH3 | R98-Y105 |
| CDRL1 | A25-L33 |
| CDRL2 | Y49-R53 |
| CDRL3 | Q90-W96 |

8TQ9

| CDR | S19.8 |
|---|---|
| CDRH1 | S25-Y33 |
| CDRH2 | N52-G56 |
| CDRH3 | R99-F107 |
| CDRL1 | T25-N32 |
| CDRL2 | S49-T53 |
| CDRL3 | Q90-L96 |

Supplementary Figure 5. Fab Sequence Alignments and CDR ranges. Amino acid sequences of the indicated Fab H and L chains, as deduced from their respective nucleotide sequences and confirmed structurally were aligned with Clustal Omega https://www.ebi.ac.uk/seqdb/confluence/display/JDSAT/Clustal+Omega+Help+and+Documentation (*91*). These are shown for are shown for the $V_H$ (A) and $V_L$ (B) regions of the indicated antibodies. Dashes indicate amino acid residues that were not observed in electron density maps. Color highlights indicate structurally defined loops corresponding to the CDRs tabulated in panel C.

| Fab | Chains | MHC Residue (chain A) | Ab Residue (H) | Distance (Å) |
|---|---|---|---|---|
| **34-5-8** | AH | GLU104 | SER102 | 3.18 |
| | AH | ASP106 | ASN52 | 2.41 |
| | AH | GLY107 | THR30 | 3.71 |
| | AH | ARG108 | SER31 | 3.43 |
| | AH | LEU109 | SER31 | 2.63 |
| | AH | LEU110 | PHE101 | 3.30 |
| | AH | ARG111 | PHE101 | 4.74 |
| | AH | GLU128 | PHE101 | 3.41 |
| | AH | ASP129 | TYR100 | 3.34 |
| | AH | LEU130 | TYR32 | 3.64 |
| | AH | LYS131 | TYR100 | 3.38 |
| | AH | GLU154 | SER1 | 4.95 |
| | AH | ARG157 | GLY26 | 4.27 |
| | AH | GLU161 | SER31 | 2.57 |
| | AH | GLY162 | ALA28 | 4.27 |
| | AH | VAL165 | SER31 | 3.92 |
| | AH | ARG169 | TYR54 | 3.33 |
| | | **MHC Residue (chain A)** | **Ab Residue (L)** | |
| | AL | ASP106 | GLU97 | 4.90 |
| | AL | ARG108 | GLU97 | 3.10 |
| | AL | ASN127 | TYR53 | 4.37 |
| | AL | GLU128 | ARG54 | 2.76 |
| | AL | ASP129 | SER60 | 4.03 |
| | AL | THR132 | SER60 | 3.27 |

| Fab | Chains | MHC Residue (chain A) | Ab Residue (H) | Distance (Å) |
|---|---|---|---|---|
| **34-2-12** | AH | ARG194 | THR96 | 4.92 |
| | AH | GLU196 | TRP99 | 2.57 |
| | AH | GLY197 | ALA97 | 3.29 |
| | AH | ASP198 | THR96 | 2.81 |
| | AH | LEU219 | ARG70 | 2.83 |
| | AH | ASN220 | ARG70 | 3.41 |
| | AH | GLY221 | ARG70 | 4.47 |
| | AH | GLU222 | THR24 | 2.38 |
| | AH | LEU224 | ASN27 | 4.20 |
| | AH | GLN226 | GLY22 | 3.29 |
| | AH | GLU227 | ASN27 | 2.84 |
| | AH | VAL247 | ASN27 | 4.18 |
| | AH | VAL248 | ASN27 | 2.72 |
| | AH | VAL249 | ASN27 | 3.51 |
| | AH | PRO250 | THR96 | 3.24 |
| | AH | LEU251 | THR96 | 4.28 |
| | AH | LYS253 | ASP51 | 2.62 |
| | AH | LYS256 | ASP51 | 3.73 |
| | AH | TYR257 | ASN27 | 2.29 |
| | | **MHC Residue (chain A)** | **Ab Residue (L)** | |
| | AL | PRO195 | PHE45 | 3.82 |
| | AL | GLU196 | PHE45 | 3.66 |

| Fab | Chains | MHC Residue (chain A) | Ab Residue (H) | Distance (Å) |
|---|---|---|---|---|
| **28-14-8** | AH | ARG111 | LYS19 | 4.79 |
| | AH | GLU128 | LYS19 | 3.86 |
| | AH | ASP212 | TYR57 | 3.22 |
| | AH | ILE213 | ASN52 | 4.79 |
| | AH | THR214 | TRP33 | 3.26 |
| | AH | THR216 | PHE100 | 3.78 |
| | AH | GLY221 | TYR101 | 2.64 |
| | AH | GLU221 | TYR102 | 4.11 |
| | AH | GLU223 | TYR102 | 3.16 |
| | AH | THR225 | THR31 | 3.27 |
| | AH | GLN226 | TYR32 | 3.21 |
| | AH | ARG260 | TYR101 | 4.78 |
| | AH | TYR262 | TRP33 | 3.47 |

| | | | | |
|---|---|---|---|---|
| | AH | GLU264 | TYR57 | 3.13 |
| | | **MHC Residue (chain B)** | **Ab Residue (H)** | **Distance (Å)** |
| | BH | ASP59 | LYS74 | 4.87 |
| | | **MHC Residue (chain A)** | **Ab Residue (L)** | **Distance (Å)** |
| | AL | GLN218 | TYR32 | 4.82 |
| | AL | LEU219 | ARG53 | 3.14 |
| | AL | ASN220 | ARG53 | 3.09 |
| | AL | GLY221 | TYR49 | 3.22 |
| | AL | GLU222 | TYR49 | 3.79 |
| | AL | ARG260 | GLY91 | 3.04 |
| | AL | TYR262 | PHE94 | 3.80 |
| | AL | PRO267 | THR93 | 3.57 |
| | AL | GLU268 | ASN92 | 3.68 |
| | AL | PRO269 | ASN92 | 3.16 |
| | AL | THR271 | TYR32 | 3.12 |
| | AL | PHE17 | THR49 | 3.96 |

| **Fab** | **Chains** | **MHC Residue (chain A)** | **Ab Residue (H)** | **Distance (Å)** |
|---|---|---|---|---|
| **S19.8** | AH | SER13 | TYR100 | 4.67 |
| | AH | ARG14 | TYR100 | 3.79 |
| | AH | PRO15 | TYR100 | 4.14 |
| | AH | PHE17 | TYR100 | 4.16 |
| | AH | GLY91 | TYR99 | 3.87 |
| | AH | SER92 | TYR99 | 3.62 |
| | | **MHC Residue (chain B)** | **Ab Residue (H)** | |
| | BH | TYR-1 | ASP31 | 3.32 |
| | BH | ALA0 | ASP31 | 3.42 |
| | BH | ILE1 | TYR54 | 3.64 |
| | BH | GLN2 | TYR54 | 3.11 |
| | BH | THR4 | ASN55 | 4.94 |
| | BH | PRO32 | TYR102 | 4.40 |
| | BH | HIS34 | TYR102 | 3.48 |
| | BH | GLU36 | PRO104 | 3.91 |
| | BH | VAL82 | TYR106 | 4.90 |
| | BH | LYS83 | PRO104 | 3.24 |
| | BH | HIS84 | PRO104 | 4.01 |
| | BH | ALA85 | TYR102 | 3.49 |
| | BH | SER86 | TYR33 | 3.66 |
| | BH | MET87 | ARG99 | 2.92 |
| | BH | ALA88 | TYR33 | 2.96 |
| | BH | GLU89 | TYR106 | 3.36 |
| | BH | PRO90 | TYR106 | 3.25 |
| | | **MHC Residue (chain A)** | **Ab Residue (L)** | |
| | AL | PHE17 | THR49 | 3.976 |
| | | **MHC Residue (chain B)** | **Ab Residue (L)** | |
| | BL | GLU36 | ASN28 | 4.66 |
| | BL | GLN38 | ASP26 | 3.32 |
| | BL | LYS45 | ASP26 | 2.40 |
| | BL | ARG81 | ASP89 | 3.14 |
| | BL | LYS83 | SER87 | 3.74 |
| | BL | ALA88 | MET90 | 3.75 |
| | BL | GLU89 | MET90 | 3.54 |
| | BL | PRO90 | ASP88 | 3.74 |

Supplementary Table 1. Contacts between the indicated Fab and H2-$D^d$ (for 34-5-8, 34-2-12, and S19.8) or H2-$D^b$ (for 28-14-8). Contacting residues with distances $\leq$ 5 Å were evaluated as described in Methods. “A” refers to the indicated MHC-I heavy chain, “B” to indicated MHC-I L chain ($b_2$m), “H” and “L” to Fab heavy and light chains respectively.

| Complex | PDB | Chains | BSA-H | BSA-L | BSA-(H+L) |
|---|---|---|---|---|---|
| Fab34-5-8/$D^d$_A | 8TQ8 | A (H,L) | 714 | 211 | 925 |
| Fab34-5-8/$D^d$_B | 8TQ8 | B (H,L) | 0 | 0 | 0 |
| Fab34-2-12/$D^d$_A | 8TQ7 | A (H,L) | 679 | 78 | 757 |
| Fab34-2-12/$D^d$_B | 8TQ7 | B (H,L) | 0 | 0 | 0 |
| Fab28-14-8/$D^b$_A | 8TQA | A (H,L) | 503 | 349 | 852 |
| Fab28-14-8/$D^b$_B | 8TQA | B (H,L) | 11 | 0 | 11 |
| FabS19.8/$D^d$_A | 8TQ9 | A (H,L) | 139 | 52 | 191 |
| FabS19.8/$D^d$_B | 8TQ9 | B (H,L) | 663 | 243 | 906 |

Supplementary Table 2. Buried surface area (BSA) at interface between Fab and pMHC. Buried surface areas for each of the indicated interfaces were calculated with PDBePISA (*89*). Values are in Å$^2$.

| **Fab** | X-ray Model (PDB) | AF-M Model | X-ray Elbow Angle (°) | AF-M Elbow Angle (°) | **Δ Elbow Angle (°)** | Limit-l | Limit-h |
|---|---|---|---|---|---|---|---|
| **34-5-8** | 8TQ8 | 8TQ8-AF | 150 | 136 | **14.0** | 113 | 119 |
| **34-2-12** | 8TQ7 | 8TQ7-AF | 135 | 136 | **1.0** | 109 | 117 |
| **28-14-8** | 8TQA | 8TQAAF | 173 | 145 | **28.0** | 107 | 116 |
| **S19.8** | 8TQ9 | 8TQ9AF | 152 | 168 | **16.0** | 107 | 116 |

Supplementary Table 3. Comparison of elbow angles of X-ray and AF-M structural models. "Limit_l" and "limit_h" indicate the residues defining the ends of variable domain of the light chain and the heavy chain respectively. The measurement of elbow angle is calculated by using the Python Script of "elbow_angle.py" (*72*) imported to PyMol.

| **Molecular Complex** | **PDB for comparison** | **DockQ score** | **Evaluation** |
|---|---|---|---|
| H2-D$^d$/$\beta_2$m | pH2-D$^d$ to $\beta_2$m from 8TQ8 (complex with 34-5-8) | 0.448 | Acceptable |
| H2-D$^d$/$\beta_2$m | pH2-D$^d$ to $\beta_2$m from 8TQ7 (complex with 34-2-12) | 0.850 | High |
| H2-D$^b$/$\beta_2$m | pH2-D$^d$ to $\beta_2$m from 8TQA (complex with 28-14-8) | 0.319 | Acceptable |
| H2-D$^d$/$\beta_2$m | pH2-D$^d$ to $\beta_2$m from 8TQ9 (complex with S19.8) | 0.474 | Acceptable |
| Fab34-5-8 H to L | Fab H to L from 8TQ8 | 0.525 | Acceptable |
| Fab34-2-12 H to L | Fab H to L from 8TQ7 | 0.879 | High |
| Fab28-14-8 H to L | Fab H to L from 8TQA | 0.606 | Medium |
| FabS19.8 H to L | Fab H to L from 8TQ9 | 0.718 | Medium |
| Fab34-5-8/H2-D$^d$ | Complete Fab to pMHC of 8TQ8 | 0.027 | Incorrect |
| Fab34-2-12/H2-D$^d$ | Complete Fab to pMHC of 8TQ7 | 0.007 | Incorrect |
| Fab28-14-8/H2-D$^b$ | Complete Fab to pMHC of 8TQA | 0.015 | Incorrect |
| FabS19.8/H2-D$^d$ | Complete Fab to pMHC of 8TQ9 | 0.023 | Incorrect |

Supplementary Table 4. Comparison of docking of X-ray and AF-M complexes using DockQ. To compare the veracity of pMHC/b$_2$m complexes, Fab H/L complexes, and docked Fab/pMHC-I complexes, the X-ray and AF-M coordinates were compared using DockQ (*56*). For the indicated complexes a value of DockQ was calculated. As suggested, DockQ scores were considered: incorrect, 0<0.2; acceptable, 0.2<0.5; medium, 0.5<0.8; and high, 0.8<1.0.
